# The nucleoporin Nup170 mediates subtelomeric gene silencing through the Ctf18-RFC complex and PCNA

**DOI:** 10.1101/2022.06.17.496627

**Authors:** Sanjeev Kumar, Maxwell L. Neal, Song Li, Arti T. Navare, Fred D. Mast, Michael P. Rout, John D. Aitchison

## Abstract

The nuclear pore complex (NPC) physically interacts with chromatin and regulates gene expression. The inner ring nucleoporin Nup170 has been implicated in chromatin organization and the maintenance of gene silencing in subtelomeric regions. To gain insight into how Nup170 regulates this process, we used protein-protein interaction, genetic interaction, and transcriptome correlation analyses to identify the Ctf18-RFC complex, an alternative proliferating cell nuclear antigen (PCNA) loader, as a facilitator of the gene regulatory functions of Nup170. The Ctf18-RFC complex is recruited to a subpopulation of NPCs that lack the nuclear basket proteins Mlp1 and Mlp2. In the absence of Nup170, PCNA levels on DNA are reduced, resulting in the loss of silencing of subtelomeric genes. Increasing PCNA levels on DNA by removing Elg1, which is required for PCNA unloading, rescues subtelomeric silencing defects in *nup170*Δ. The NPC therefore mediates subtelomeric gene silencing by regulating PCNA levels on DNA.

## INTRODUCTION

Nuclear pore complexes (NPCs) are large proteinaceous assemblies embedded in the nuclear envelope (NE) that serve as the only conduits for nucleocytoplasmic transport (Aitchison and Rout, 2012). The yeast NPC, an over five-hundred protein megacomplex, is made up of an assembly ∼30 different proteins, termed nucleoporins (Nups) (Alber et al., 2007; Kim et al., 2018), that are each present in multiple copies per NPC. These nucleoporins assemble to form higher-order modular structures called spokes, and eight coaxially arranged spokes form a symmetrical, cylindrical channel. Each spoke is composed of different sub-complexes arranged to form the outer, inner, and membrane rings, cytoplasmic filaments, and a nuclear basket (Alber et al., 2007; Kim et al., 2018). The central channel of the NPC is composed of nucleoporins rich in disordered Phe-Gly (FG) repeat motifs that line the pore and that confer selectivity to transport of cargos between the nucleus and cytoplasm (Aitchison and Rout, 2012).

NPCs also function as positional beacons or scaffolds in the NE with various roles in chromatin organization and gene regulation. Thus, they influence heterochromatin states, nucleosome organization, chromatin boundaries and telomere localization. This influence, in turn, leads to the regulation of transcriptional activation, transcriptional repression, transcriptional memory, and subtelomeric silencing (Ptak et al., 2014; Van de Vosse et al., 2013; Dilworth et al., 2001; Kadota et al., 2020; Light et al., 2010; Galy et al., 2000).

During S-phase, newly synthesized DNA is packaged into nucleosomes by replication-coupled nucleosome assembly, which prevents DNA damage and preserves the integrity of the genome (Serra-Cardona and Zhang, 2018). Replication forks act as organizing centers for a host of chromatin modifiers that are required to reassemble chromatin after replication. Impairment of replication fork progression during DNA replication by endogenous or exogenous factors can result in DNA damage and trigger DNA damage response pathways. Some of the stalled forks become spatially segregated to the nuclear periphery and the NPC has been implicated in the resolution of these stalled forks and in restarting DNA replication (Lamm et al., 2021; Whalen and Freudenreich, 2020). However, the precise role that the NPCs play in this process remains poorly understood.

The inner ring nucleoporin Nup170 interacts with chromatin, and it is implicated in the maintenance of heterochromatin and genome stability (Kerscher et al., 2001; Van de Vosse et al., 2013). Nup170 is required for telomere tethering to the NE, and disruption of Nup170 leads to altered chromatin structure and the upregulation of many genes, the majority of which are located in the otherwise silenced subtelomeric regions of chromosomes (Van de Vosse et al., 2013). Nup155, the mammalian ortholog of Nup170, also associates with, and modulates, chromatin functions, suggesting a potentially conserved role (Breuer and Ohkura, 2015; Kehat et al., 2011). Although disruption of Nup170 has profound effects on various aspects of chromatin activities, the mechanistic underpinnings of its contributions to chromatin regulation remain unclear.

To gain insight into the roles of Nup170 in chromatin organization and gene regulation, we employed molecular systems biology approaches to identify the alternative proliferating cell nuclear antigen (PCNA) loader Ctf18-RFC complex as a mediator of Nup170’s chromatin modification functions. Here, we report that the Ctf18-RFC complex physically and functionally interacts with Nup170 and PCNA levels on chromatin are reduced in cells lacking Nup170. We show loss of subtelomeric silencing in *nup170*Δ is largely rescued by increasing PCNA levels on DNA by removal of the PCNA unloader Elg1. Our results reveal a surprising new role for NPCs in the maintenance of PCNA levels on DNA and in the regulation of chromatin functions.

## RESULTS

### Systems biology approaches identify the Ctf18-RFC complex as a mediator of Nup170’s gene regulatory functions

To understand the underlying mechanism(s) of the gene regulatory functions of Nup170, we defined a protein-protein interaction (PPI) map of transient and indirect vicinal interactors of the NPC that are centered around Nup170. A chimeric Nup170-GFP fusion protein was affinity purified from whole cell extracts prepared under non-stringent isolation conditions from cryo-milled yeast and its co-enriching interacting proteins were identified by mass spectrometry (Fig. S1A) (Hakhverdyan et al., 2015). This analysis revealed 557 non-nucleoporin interacting proteins of Nup170, of which 210 have previously been localized to the nucleus and therefore represent putative candidate facilitators of Nup170’s role in regulating chromatin (Fig. 1A). To identify proteins with putative functions related to Nup170’s role in gene expression, we parsed this list through a rubric of: 1) Functional relevance revealed by a genetic interaction with Nup170; and 2) Among these genetically interacting genes, the requirement that its deletion mutant phenocopy *nup170*Δ as measured by gene expression profiles. There are 415 reported genetic interactors of Nup170 in BioGRID (https://thebiogrid.org/, Stark et al., 2006), and 33 of these genetic interactors were also identified as vicinal interactors of Nup170 (Fig S1A), suggesting some of these proteins may function with, or redundantly to, Nup170. We assessed the correlation of gene expression profiles using a compendium of transcriptomic profiles of ∼1,500 individual deletion strains (Kemmeren et al., 2014) called the “deleteome”. This transcriptome correlation analysis identified 40 individual gene deletions (Fig. 1A) whose expression profiles were significantly correlated with the *nup170Δ* strain. Among genetic and physically interacting proteins, the deletions with significant transcriptome correlation scores were chromosome transmission fidelity 18 (Ctf18) (Mayer et al., 2001) and ubiquitin-specific protease 3 (Ubp3)(Baker et al., 1992). We prioritized Ctf18 for further study based on its direct and defined roles in chromatin organization, telomere maintenance and telomere recruitment to the nuclear periphery (Gellon et al., 2011; Hiraga et al., 2006; Kubota et al., 2011; Liu et al., 2020; Ogiwara et al., 2007; Stokes et al., 2020).

**Figure 1.**
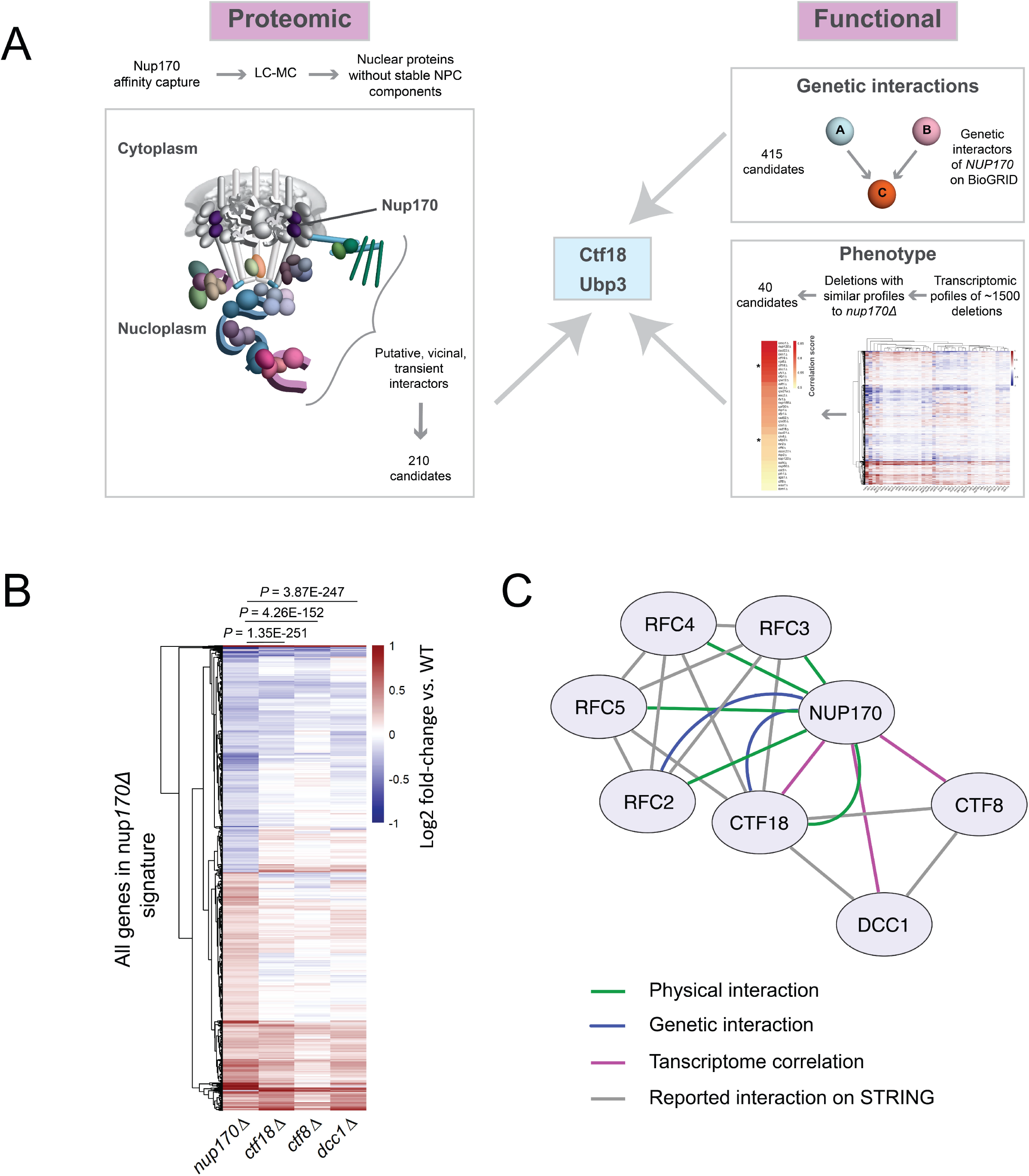
Systems biology approaches identify the Ctf18-RFC complex as a mediator of Nup170’s gene regulatory functions. (A) Schematic of our approach to identify and prioritize candidates that mediate gene regulatory roles of the nucleoporin Nup170. The LC-MS analysis of affinity-purified Nup170-GFP identified 210 vicinal interactors on the nucleoplasmic side, 33 of which have known genetic interactions with *NUP170*. Among genetically and physically interacting candidates, transcriptomic profiles of *ctf18*Δ and *ubp3*Δ strains were significantly similar to the *nup170*Δ strain in a compendium of deletion strain transcriptomes. (B) Heatmap showing gene expression profiles of the *nup170*Δ strain and the three (*ctf18*Δ, *ctf8*Δ and *dcc1*Δ) viable deletion strains for Ctf18-RFC complex constituents. (C) Cytoscape network illustrating interaction between Nup170 and the Ctf18-RFC complex.

Ctf18 is a subunit of the highly conserved heptameric alternative DNA sliding clamp loader replication factor C (RFC) complex (Mayer et al., 2001). The canonical RFC complex consists of five essential proteins of the AAA+ ATPase family containing one large (Rfc1) and four small (Rfc2-5) subunits (Bowman et al., 2004; Yao and O’Donnell, 2012). In the alternative clamp loader Ctf18-RFC complex, Rfc1 is replaced by Ctf18 and its C-terminus is bound to two additional proteins Ctf8 and defective in sister chromatid cohesion 1 (Dcc1) (Grabarczyk et al., 2018). Consistent with a functional relationship between the Ctf18-RFC complex and Nup170, each of the four small subunits - Rfc2, Rfc3, Rfc4 and Rfc5 - were present in our Nup170 vicinal interactome dataset (Fig. 1C, Table S1). Rfc2-5 are essential proteins, and therefore transcriptome profiles cannot be obtained for their deletions. However, we found expression profiles for deletion strains of the two non-essential components *DCC1* and *CTF8* were significantly correlated with that of *nup170*Δ, and *ctf18*Δ (Fig. 1A & B).

To confirm the physical interaction between Nup170 and the Ctf18-RFC complex we performed reciprocal affinity capture of Ctf18-GFP and Rfc3-GFP under different extraction conditions and probed the eluates for Nup170-3FLAG by immunoblot. Consistent with our affinity purification mass spectrometry data, we found that Nup170-3FLAG co-purified with both Ctf18-GFP and Rfc3-GFP (Fig. S1B) confirming the likely vicinal association between the Ctf18-RFC complex and Nup170.

### The Ctf18-RFC complex is recruited to a subset of NPCs lacking the nuclear basket nucleoporins Mlp1 and Mlp2

Accumulating evidence suggests that structurally heterogeneous NPCs exist within a single cell, which may in turn facilitate a variety of functions (Fernandez-Martinez and Rout, 2021). Recent work by Akey et al., (2022) suggests that at least three major structural forms of NPCs exist in a single NE, which may aid in functional specialization of the nuclear periphery. The three isoforms of the NPC contain either a single or double outer ring on their nucleoplasmic side, or lack the nuclear basket. Separately, Nup170 has been shown to be a constituent of a distinct ‘Snup’ complex of nucleoporins that interacts with the silencing factor sirtuin 4 (Sir4) to recruit subtelomeres to the nuclear periphery (Lapetina et al., 2017). The Snup complex is physically distinct from mature NPCs, and lacks the flexible-connector Nups of the inner ring complex (Nup53 and Nup59), and also the nuclear basket Nups (Nup60, Nup1, and Mlp1). Given this background, we asked two questions: 1) is the Ctf18-RFC complex recruited to all NPCs? and if not; 2) can we define the composition of the NPC isoform(s) with which it interacts? To answer the first question, we used a split GFP system (Cabantous and Waldo, 2006) where two proteins of interest are separately tagged with two non-fluorescent parts of GFP (GFP_1-10_ and GFP_11_). This system has been used to study in vivo, dynamic, and sub-stoichiometric protein-protein interactions (Hu and Kerppola, 2003, 2003; Hu et al., 2002). If the two proteins physically interact, the two GFP fragments are brought into close proximity, assemble, and reconstitute fluorescence (Hu et al., 2002; Smoyer et al., 2016). Using Nup170 as bait, we tested each component of the Ctf18-RFC complex and observed a few GFP puncta per cell that localized to the NE and overlapped with a portion of NPCs labelled with Nup188-mCherry (Fig. 2A). These results indicate a close physical interaction between Nup170 and the Ctf18-RFC complex in vivo, and that the Ctf18-RFC complex is recruited to only a small subset of NPCs (Fig. 2A). These findings are likely specific for a subset of the total Ctf18-RFC complex because despite the high abundance of some members, e.g., Rfc3 has ∼5000 copies per cell (Ho et al., 2018), only a few GFP foci were observed. As a control, we did not observe GFP puncta when Rfc3 was tested in combination with the FG-containing Nup49 (Wente et al., 1992), suggesting that not all Nups are accessible, and in particular, Nup49 of the central core of the NPC was inaccessible to the Ctf18-RFC complex.

**Figure 2.**
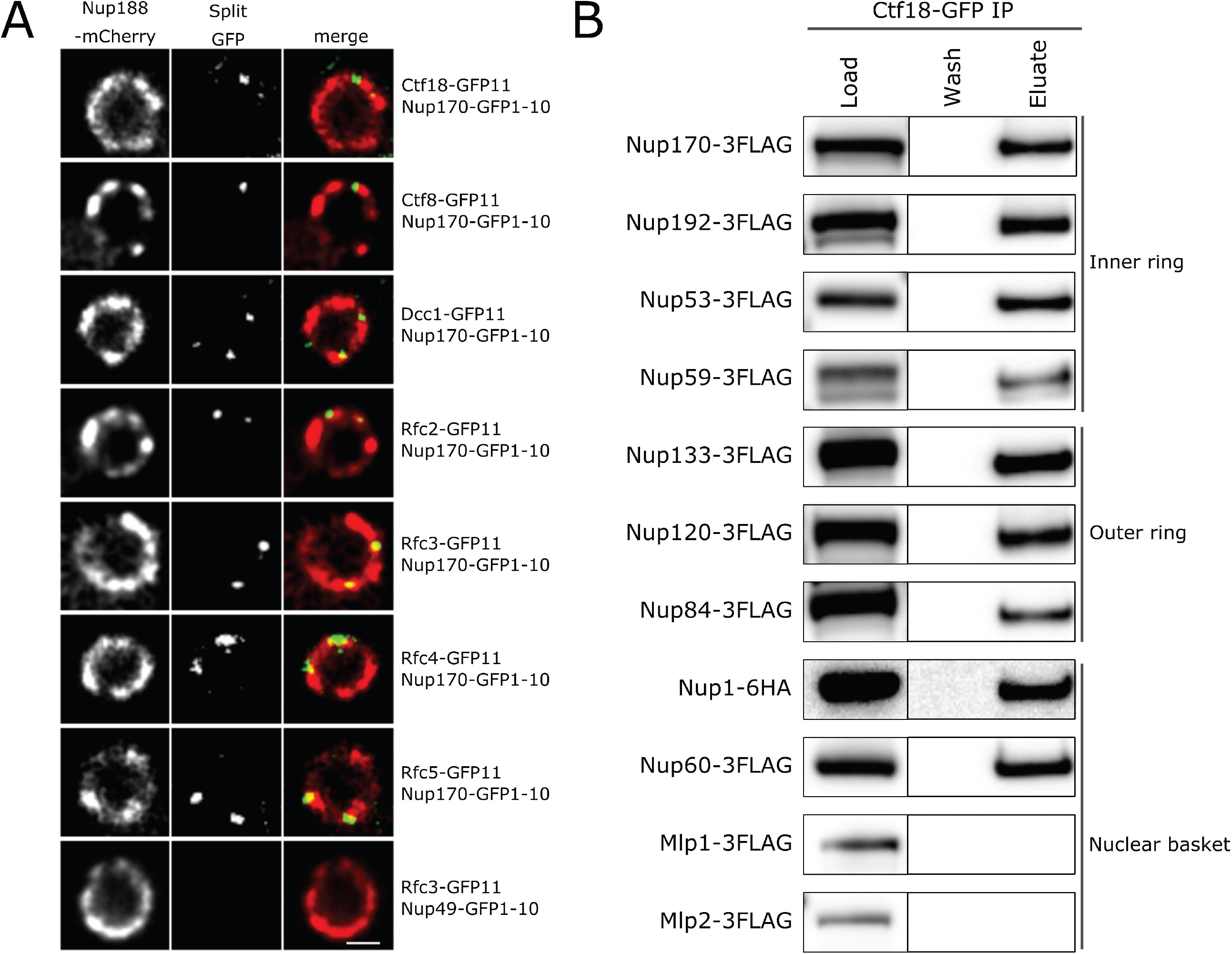
Ctf18-RFC is recruited to a subset of NPCs lacking the nuclear basket nucleoporins Mlp1 and Mlp2. (A) Images of the nuclei of cells co-expressing Nup170-GFP1-10 and GFP11 tagged components of the Ctf18-RFC complex. The bottom panel image shows a cell co-expressing Nup49-GFP1-10 and Rfc3-11. In these cells the NPCs are marked by Nup188-mCherry. A single plane image from the acquired z-stack is shown. Bar = 1 µm (B) Ctf18-GFP fusion protein was affinity purified from cell lysates containing 3FLAG-tagged nucleoporins or Nup1-6HA. Eluates were analyzed by immunoblotting using anti-FLAG or anti-HA antibodies to detect indicated nucleoporins.

To assay for the NPC isoform(s) associating with the Ctf18-RFC complex, we affinity purified Ctf18 and probed the eluate for a panel of nucleoporins. In addition to Nup170, we tested for members of the inner ring complex (Nup192, Nup53 and Nup59), the outer ring complex (Nup133, Nup120 and Nup84), and the nuclear basket (Nup1, Nup60, Mlp1 and Mlp2), but not the cytoplasmic Nups. All tested inner ring, outer ring, and nucleoplasmic FG Nups copurified with Ctf18-GFP (Fig. 2A). However, under the condition used for the affinity purification, we did not detect a physical association between Ctf18-GFP and the nuclear basket proteins Mlp1 and Mlp2. Taken together, these results confirm a physical association between Ctf18-RFC and a subset of NPCs that are either distinct from or not limited to Snup complexes, and that are devoid of the major basket proteins Mlp1 and Mlp2 (Niepel et al., 2005; Strambio-de-Castillia et al., 1999).

### Ctf18 interaction with the NPC is enriched in S phase

Chromatin immunoprecipitation analyses have shown that the Ctf18-RFC complex localizes to replication forks and is bound to chromatin throughout S phase (Crabbé et al., 2010; Lengronne et al., 2006; Liu et al., 2020). To test if the observed Ctf18-RFC interaction with NPCs differed through the cell cycle, we analyzed the localization of Ctf18-GFP during various stages of the cell cycle in a strain in which NPCs were marked by endogenously tagged Nup188-mCherry. Ctf18-GFP appeared mainly diffuse in the nucleoplasm during G1, but concentrated into a few foci during S phase. After completion of S phase Ctf18-GFP was redistributed and appeared diffuse in the nucleus during G2/M-phase (Fig. 3A, B). We quantified the localization of Ctf18-GFP foci relative to Nup188-mCherry in individual cells and found that the majority of them were proximal to the NPC and/or overlapping with Nup188-mCherry signal (Fig 3C). To further evaluate cell cycle dependent interactions between Ctf18 and the NPC, we affinity purified Ctf18-GFP from cells harvested at different timepoints after release from G1 arrest and probed eluates for Nup170-3FLAG by immunoblotting. Consistent with our microscopy data, we found that the interaction between Ctf18 and Nup170 was highest during S-phase (Fig. 3D).

**Figure 3.**
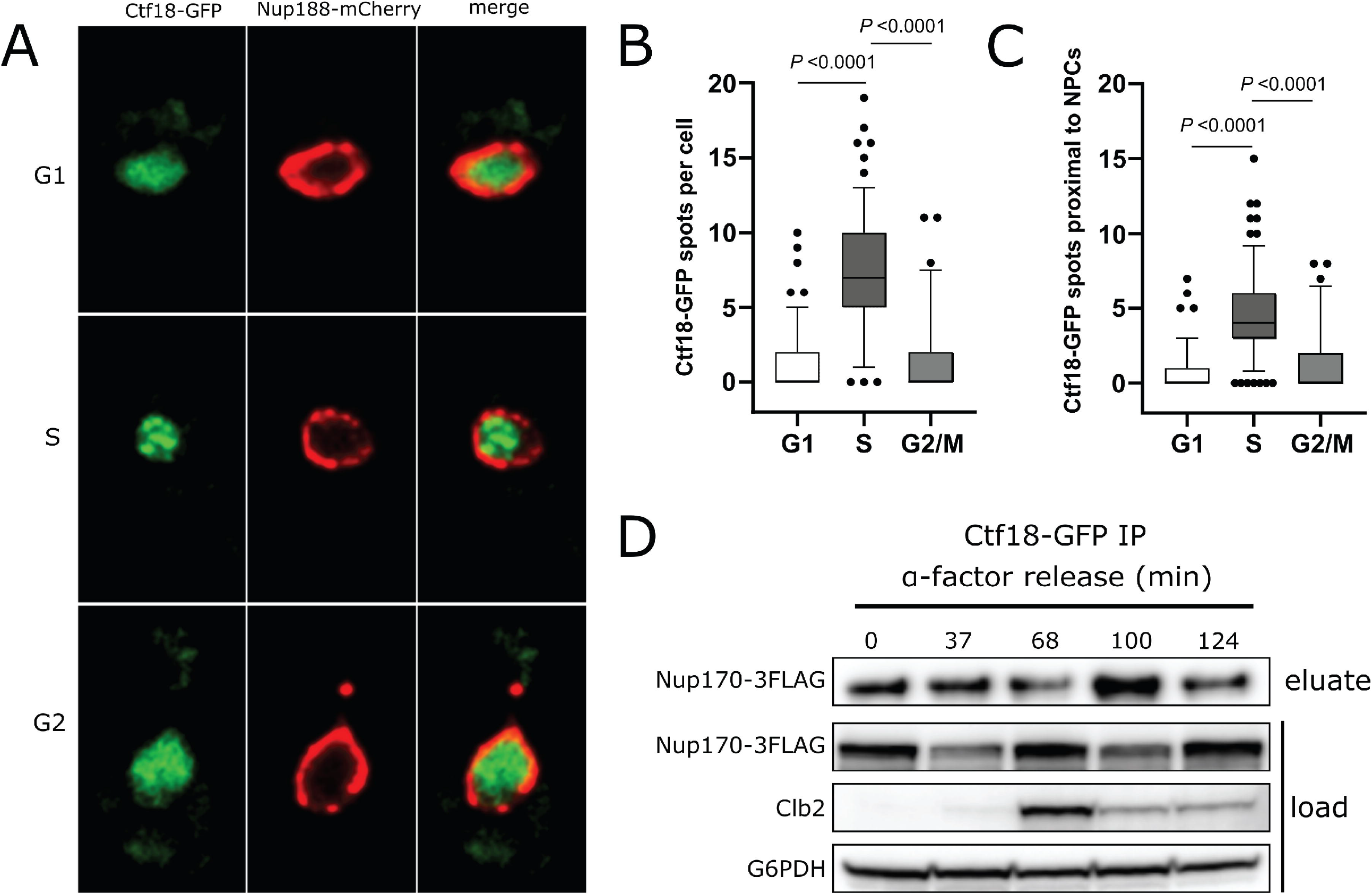
Ctf18 interaction with the NPC is maximal during S-phase. (A) Cells expressing genomically encoded Nup188-mCherry and Ctf18-GFP were arrested in G1 phase by α-factor and images were acquired at regular intervals for 110 min after release from arrest. Representative images are shown for different cell cycle stages. (B) Box plot showing number of Ctf18-GFP spots per cell at the G1, S and G2/M cell cycle stages. (C) Box plots showing number of Ctf18-GFP spots close to NPCs at those stages. The *P* values in C and D are from Student’s t-test. (D) Cells producing Ctf18-GFP and Nup170-3FLAG were arrested in G1 and samples were collected at indicated timepoints after release. The Ctf18-GFP fusion protein was affinity purified from cell lysates and the eluate was analyzed by immunoblotting to detect Nup170-3FLAG. The Clb2 levels in cell lysates (load) are indicative of cell cycle stage. Anti-FLAG, Clb2 and G6PHD antibodies were used for immunoblotting.

### Nup170 functions in subtelomeric gene silencing and DNA repair, but does not affect sister chromatid cohesion

Nup170 plays a role in telomere positioning to the nuclear periphery and controls subtelomeric silencing (Van de Vosse et al., 2013). Similar to Nup170, the Ctf18-RFC complex is required for telomere positioning to the nuclear periphery and helps facilitate silencing of a telomeric *ADE2* reporter gene (Hiraga et al., 2006; Suter et al., 2004). Indeed, the expression profiles of the *NUP170* deletion strain and deletion strains of the three non-essential components of Ctf18-RFC complex-*ctf18*Δ, *ctf8*Δ, and *dcc1*Δ-were all highly enriched for upregulated subtelomeric genes (hypergeometric test, FDR-adjusted *P*-values < 0.05) (Fig. 4A). These results further confirm the roles of Nup170 and the Ctf18-RFC complex in the regulation of subtelomeric gene silencing.

**Figure 4.**
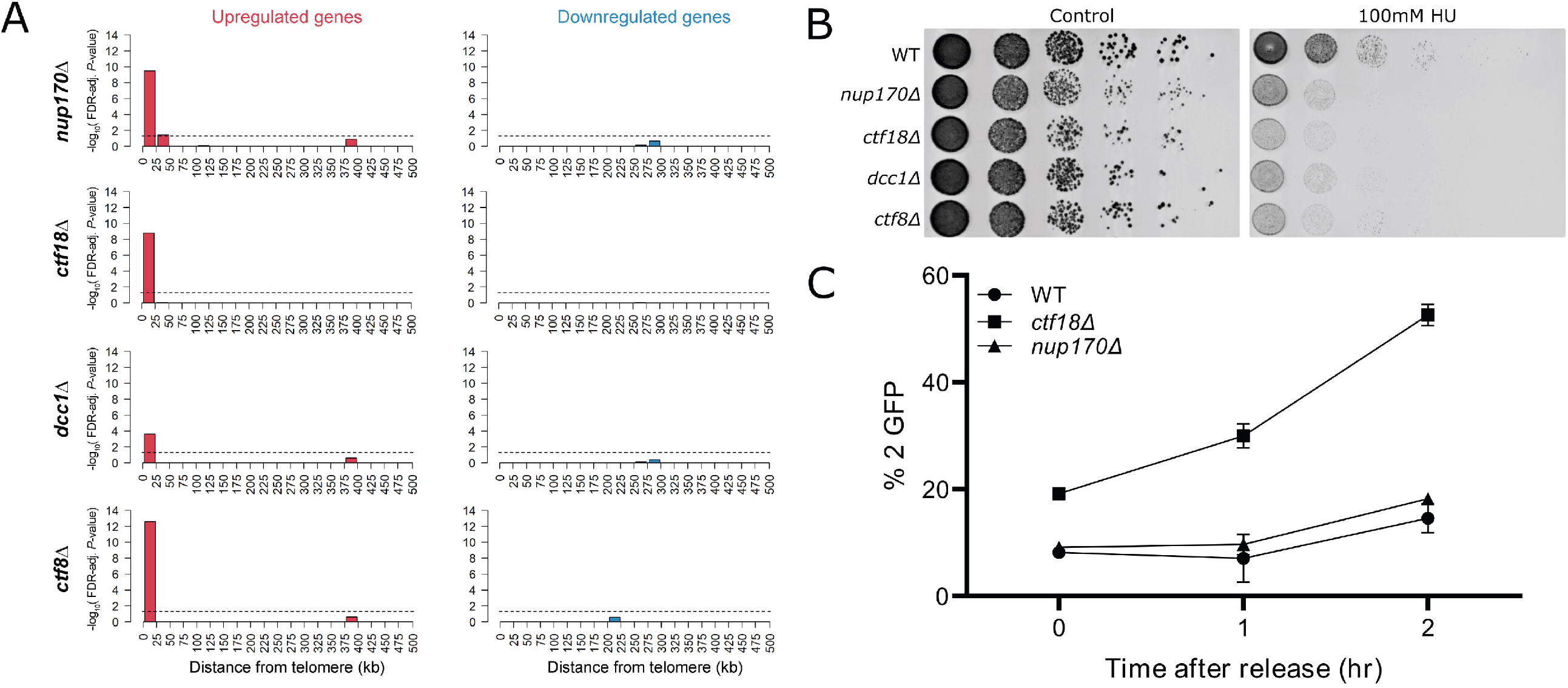
Nup170 functions in subtelomeric silencing and replication checkpoint but does not affect sister chromatid cohesion. (A) Enrichment of differentially expressed genes in the *nup170*Δ, *ctf18*Δ, *dcc1*Δ *and ctf8*Δ strains from the deleteome compendium according to gene distance from chromosome ends. X-axes: distance from chromosome ends in 25 kb bins. Y-axes: negative log_10_ of the enrichment test’s FDR-adjusted *P*-value. Dashed lines represent significance cutoff. (B) Log-phase cultures of the indicated strains were equalized in cellular density, serially diluted 10-fold, and spotted onto plates containing yeast extract, peptone, and dextrose (YPD) with or without 100 mM Hydroxyurea. Plates were scanned after 2 days at 30 °C. (C) Sister chromatid cohesion was assessed in three biological replicates for the indicated strains containing a lacO tandem repeat inserted at the *TRP1* locus on chromosome IV and LacI-GFP. The number of GFP spot in each cell was scored and percentages of cells with two GFP spots are plotted.

The Ctf18-RFC complex has been implicated in the replication checkpoint response pathway (Crabbé et al., 2010; Naiki et al., 2001). To test if Nup170 functions like the Ctf18-RFC complex in response to DNA damage, we evaluated the sensitivity of cells lacking Nup170 or the components of the Ctf18-RFC complex to hydroxyurea (HU), a ribonucleotide reductase inhibitor that arrests cells in S-phase and induces replication stress (Alvino et al., 2007). We found that, similar to the *ctf18*Δ, *ctf8*Δ, and *dcc1*Δ strains, cells lacking Nup170 were more sensitive to HU compared to WT (Fig. 4B). We also found that in cells treated with HU, recruitment of Ctf18 to the NPC was enhanced, suggesting its role in DNA repair (Fig. S2).

One of the main functions of the Ctf18-RFC complex is to establish sister chromatid cohesion (SCC) after DNA replication and defective SCC can lead to genomic instability (Hanna et al., 2001; Mayer et al., 2001). We investigated whether the *nup170*Δ strain also exhibited a SCC defect and whether that could explain the defect in subtelomeric silencing and the strain’s sensitivity to HU. To visualize SCC we used a strain containing lacO-repeats integrated at 12.5 kb from the centromere of chromosome IV that binds a lac repressor-GFP fusion protein (Sanchez et al., 1999). To assess SCC, cells were arrested in G1 phase using α-factor followed by release into nocodazole containing growth medium to arrest again in G2/M phase. As expected, *ctf18*Δ showed SCC defects (Fig. 4C) (Hanna et al., 2001; Mayer et al., 2001); however, SCC defects were not observed in *nup170*Δ cells. These results suggest that loss of subtelomeric silencing and the replication checkpoint defect in *nup170*Δ are not due to major defects associated with SCC.

### PCNA levels decline at stalled replication forks in *nup170*Δ

The Ctf18-RFC complex is involved in both loading and unloading PCNA onto DNA (Bermudez et al., 2003; Bylund and Burgers, 2005); however, its PCNA unloading activity is not well understood and it is unclear whether this occurs in vivo. Previous studies in *ctf18*Δ cells showed that PCNA levels decline at replication forks in cells synchronously progressing through S phase or arrested in HU (Lengronne et al., 2006; Liu et al., 2020) suggesting that Ctf18-RFC functions as the net loader. We asked whether, similar to the Ctf18-RFC complex, Nup170 was also required for maintaining PCNA levels at replication forks. To analyze PCNA binding to DNA, we synchronized WT and *nup170*Δ cells in G1 phase using α-factor followed by release into HU containing growth medium to further arrest cells in S phase. We performed PCNA ChIP-seq to visualize its chromatin binding pattern, which, expectedly, showed binding to autonomously replicating sequence (ARS) sites (Fig. 5A, Fig. S3). These results indicate that PCNA binding to ARSs was similar in WT and *nup170*Δ strains (Fig. 5A, Fig. S3); however quantitation of binding by ChIP-qPCR for four ARS sites, showed significant decreases in ARS binding in the *nup170*Δ strain compared to WT cells (Fig. 5B).

**Figure 5.**
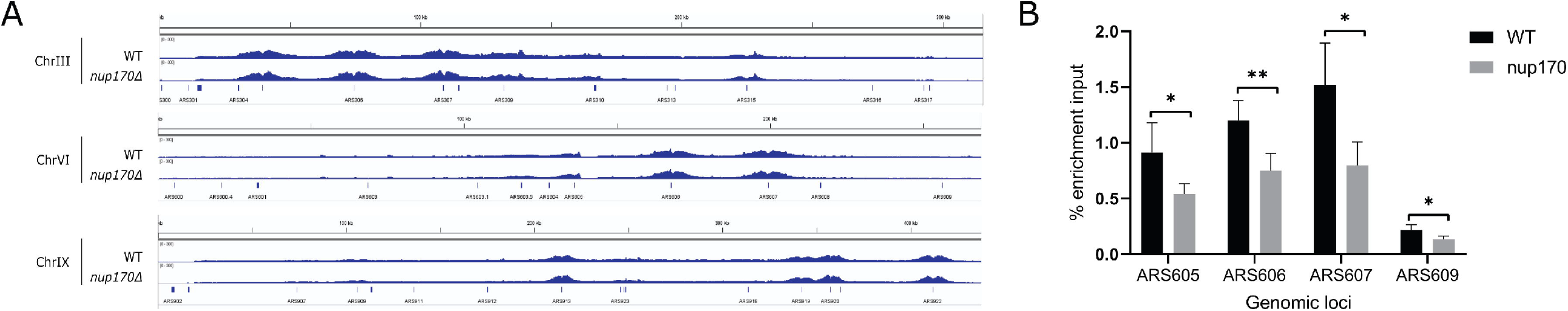
PCNA levels on chromatin declines in *nup170Δ*. (A) Cells were synchronized in G1 and released into HU-containing YPD medium to arrest them in S phase, and ChIP was performed using an anti-PCNA antibody. Normalized PCNA binding profiles along chromosomes III, VI and IX obtained by ChIP-seq analysis visualized with the Integrative Genomics Viewer. (B) PCNA enrichment at the indicated ARS sites was analyzed by qPCR. Means ± SD from four experiments are shown, \**P* < 0.05, \*\**p* < 0.01. The *P* is from Student’s t-test.

### Deleting *ELG1* in *nup170Δ* rescues subtelomeric silencing

We reasoned that if the subtelomeric silencing defect in *nup170*Δ cells stems from the reduction of PCNA levels on DNA, then increasing the PCNA levels on DNA would reverse this defect. Enhanced level of genomic instability 1 (Elg1)-RFC complex is an alternative RFC complex for unloading PCNA from chromatin, and its removal elevates PCNA levels on chromatin (Bellaoui et al., 2003; Kubota et al., 2013). Consistent with our hypothesis, analysis of the deleteome compendium data reveals that loss of Elg1 results in the decreased expression of genes significantly enriched in subtelomeric regions (Fig. 6A; Hypergeometric test, FDR-adjusted *P*-values < 0.05), essentially mirroring the increased expression observed for subtelomeric genes when Nup170 is lost (Fig 6B; Pearson’s correlation coefficient=-0.18, *P*=0.002). These data are consistent with a previous report showing that an *ELG1* deletion results in increased silencing of an *ADE2* marker gene inserted into a subtelomeric region of chromosome X (Smolikov et al., 2004).

**Figure 6.**
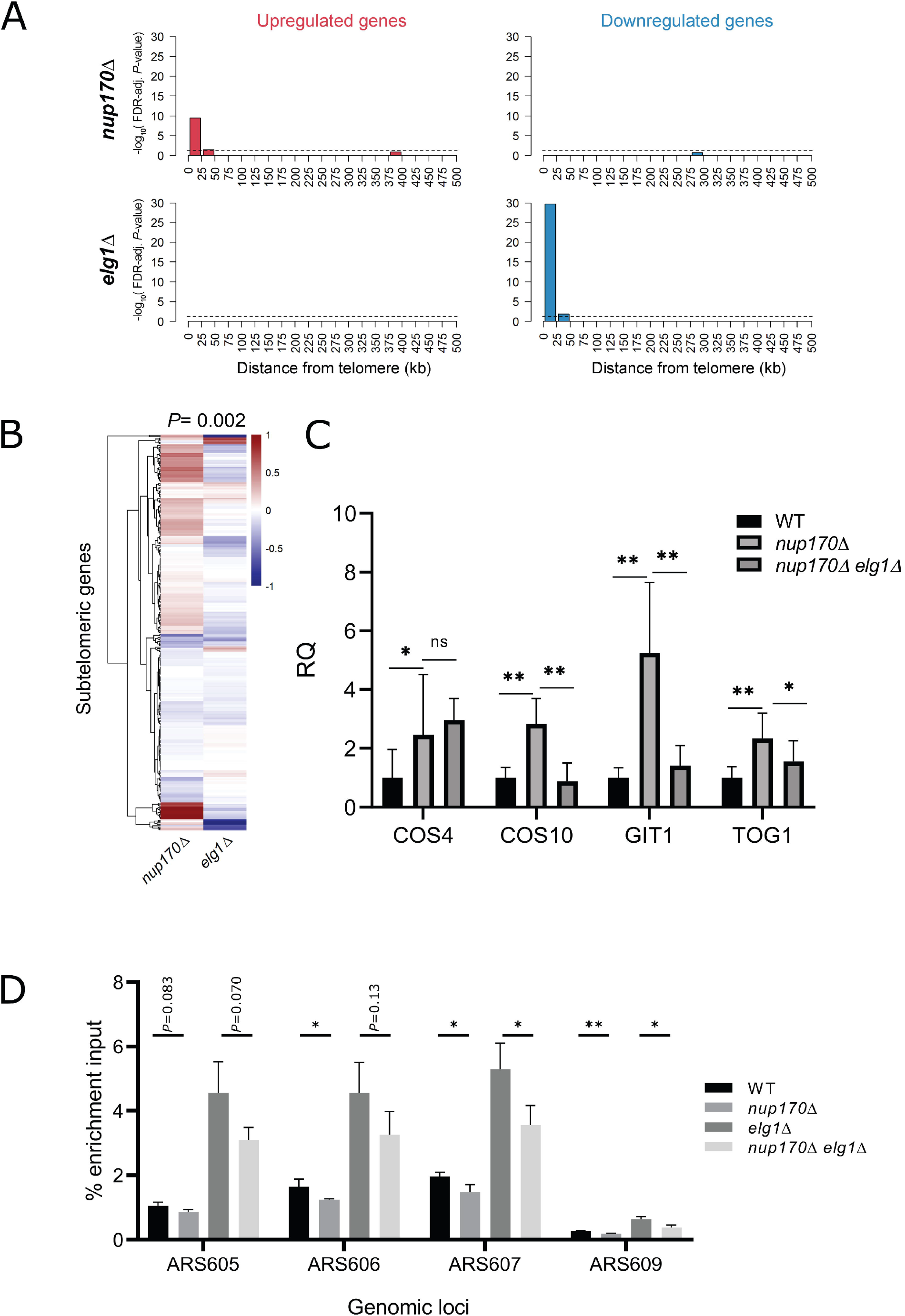
Increasing PNCA levels on DNA in *nup170*Δ rescues subtelomeric silencing defects. (A) Enrichment of differentially-expressed genes in the *nup170*Δ and *elg1*Δ strains according to gene distance from chromosome ends. X-axes: distance from telomere in 25 kb bins. Y-axes: negative log_10_ negative base 10 logarithm of the enrichment test’s FDR-adjusted *P*-value. Dashed lines represent significance cutoff. (B) Heatmap showing expression profiles of subtelomeric genes in the *nup170*Δ and *elg1*Δ strains. (C) Gene expression levels of four subtelomeric genes were measured in the WT, *nup170*Δ and *nup170*Δ *elg1*Δ strains by RT-qPCR and relative quantification (RQ) values are plotted. (D) PCNA enrichment at the indicated ARS sites was analyzed by qPCR. Means ± SD from three experiments are shown, \**P* < 0.05, \*\**p* < 0.01.

**Figure 7.**
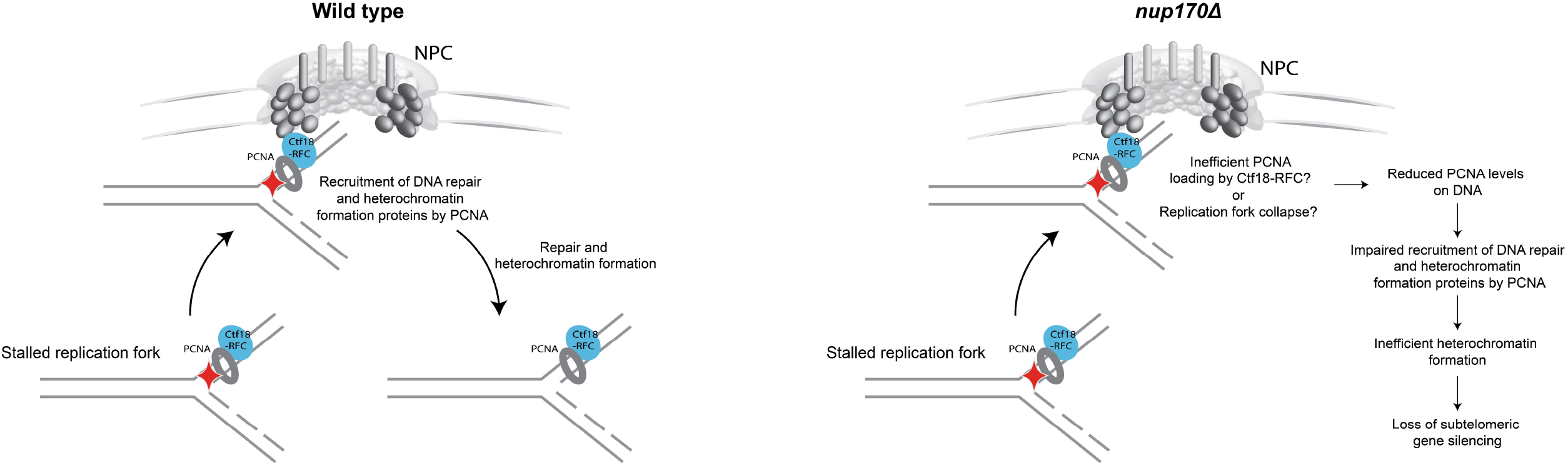
Cartoon illustrating the role of the NPC in DNA damage repair and heterochromatin formation.

To test if increasing PCNA levels on DNA in *nup170*Δ enhanced silencing, we measured the expression of a subset of subtelomeric genes by RT qPCR in *nup170*Δ and *nup170*Δ*elg1*Δ strains. *COS4, COS10, GIT1* and *TOG1* were selected based on their subtelomeric locations in the genome and previous observations that their silencing depends on Nup170 (Van de Vosse et al., 2013; Kemmeren et al., 2014). Deletion of *ELG1* in the *nup170*Δ strain reversed the effect of diminished subtelomeric silencing, with three of the four genes tested showing a significantly reduced expression in *nup170*Δ*elg1*Δ cells compared to *nup170*Δ. (Fig. 6C). Furthermore, ChIP qPCR data confirmed that PCNA levels on DNA were elevated in the *nup170*Δ*elg1*Δ strain compared to *nup170*Δ (Fig. 6D). Taken together, these results indicate that the subtelomeric silencing defect observed in *nup170*Δ cells is due to reduced PCNA levels and that increasing those levels largely restores silencing. We also investigated whether increasing PCNA levels in *nup170*Δ cells rescued the replication checkpoint defect. We found that *nup170*Δ*elg1*Δ remained susceptible to growth on HU (Fig. S4).

## DISCUSSION

Several studies in yeast and metazoans have established roles for the NPC in chromatin organization, gene regulation, and DNA repair (Ptak et al., 2014). However, in many cases the physical and functional relationship between the NPC and chromatin-modifying factors that regulate these processes remain poorly understood. The nucleoporin Nup170 plays a role in nucleosome organization and genome stability, and the loss of Nup170 leads to altered subtelomeric chromatin structure and loss of subtelomeric silencing (Kerscher et al., 2001; Van de Vosse et al., 2013). In addition to these roles, in this study we show that Nup170 also has a role in the replication checkpoint response (Fig.4B). To explore the mechanistic underpinnings of various functions of Nup170, we applied a set of comprehensive and complementary molecular systems biology approaches to identify the alternative PCNA loader Ctf18-RFC complex as a mediator of gene regulatory functions of Nup170. We show that *NUP170* deletion leads to reduced PCNA levels on DNA, which is linked to HU hypersensitivity and loss of subtelomeric gene silencing.

Evidence from the structural analyses of NPCs from several model organisms suggests significant similarities in the core scaffold regions; however, there is considerable variability in the peripheral regions of NPCs (Fernandez-Martinez and Rout, 2021). Architectural diversity in NPCs across organisms associated with differences in the stoichiometry of the structural modules, or by gain or loss of components reflects adaptations during evolution suited for each organism. To facilitate the variety of functions NPCs perform, structural heterogeneity within individual cells can be also envisioned to play a role. For example, the nuclear basket proteins Mlp1 and Mlp2 are excluded from NPCs juxtaposed to the nucleolus (Galy et al., 2004); core components of the spindle pole body coimmunoprecipitate with Mlp2, but not Mlp1 (Niepel et al., 2005); and the Snup complex has been shown to contain most nucleoporins, but lacks components of the nuclear basket (Nup60, Nup1, and Mlp1) and inner ring (Nup53 and Nup59) and interacts with Sir4 (Lapetina et al., 2017). Structural data from cryo-EM images collected from in vivo NPCs suggest three isoforms of NPCs exist within a single cell (Akey et al., 2022). These NPC isoforms contain either a single or double outer ring on the nucleoplasmic side, with an additional subpopulation of NPCs lacking a nuclear basket. Functionally, the nuclear basket nucleoporins Nup2 and Mlp1 have been shown to play a role in gene activation and transcriptional memory (Brickner et al., 2019; Tan-Wong et al., 2009), whereas Nup170 is involved in the maintenance of subtelomeric silencing and heterochromatin formation (Van de Vosse et al., 2013). We found that Ctf18 is recruited to a small subpopulation of NPCs that lack the nuclear basket proteins Mlp1 and Mlp2. Absence of these nuclear basket nucleoporins likely renders these NPCs more accessible to chromatin. These findings shed light on the functional specialization of NPCs possessing or lacking a nuclear basket. Our data are consistent with a model in which Nup170’s functional relationship with chromatin occurs via specialized NPCs lacking the nuclear basket.

Independent studies have shown that the Ctf18-RFC complex functions independently in SCC and replication checkpoint activation (Crabbé et al., 2010; Liu et al., 2020), and its SCC role is not required for telomere maintenance (Hiraga et al., 2006). The SCC defect in *ctf18*Δ was linked to reduced PCNA levels on DNA (Liu et al., 2020); however, despite reduced levels of PCNA on DNA in the *nup170*Δ strain, we did not observe significant SCC defects. These cells were susceptible to HU and had defective subtelomeric silencing suggesting that Nup170 facilitates PCNA levels on DNA upon induction of DNA damage, thus enabling DNA repair. Increased Ctf18 binding to the NPC after HU treatment further supports this conclusion. As outlined below, defects in this process can affect DNA stability, especially in heterochromatin, thereby affecting gene expression.

PCNA, which encircles DNA, is tethered to replicative DNA polymerases δ and ε, and increases their processivity (Boehm et al., 2016). In addition to its major role at replication forks during DNA replication, PCNA also serves as a loading platform for proteins involved in SCC establishment, chromatin remodeling, and the processes linked to DNA repair pathways (Mailand et al., 2013). Therefore, an intricate balance of PCNA levels on DNA that is tailored for its various activities is necessary for it to perform its functions. PCNA binds to chromatin assembly factor 1 (CAF-1) which recruits histones to accomplish DNA replication- or DNA repair-coupled nucleosome assembly (Gaillard et al., 1996; Young et al., 2020). PCNA accumulates and recruits CAF-1 to the damage site to facilitate nucleosome assembly during DNA damage repair (Essers et al., 2005; Gaillard et al., 1996; Moggs et al., 2000). Similarly, Ctf18 builds on stalled replication forks (Crabbé et al., 2010). Recently it was suggested that upon replication fork stalling Pol ε decouples from the Cdc45-MCM-GINS complex and recruits additional Ctf18-RFC complexes (Fujisawa et al., 2017), which may trigger new rounds of PCNA loading that can further recruit factors required for DNA repair and restart DNA synthesis. Expectedly, in the absence of Ctf18, PCNA levels decline on DNA (Lengronne et al., 2006; Liu et al., 2020). Here, we show that the maximum association between Ctf18 and the NPC occurs during S phase of the cell cycle (Fig. 3A, B &C); a phase that is also prone to DNA damage due to actively replicating DNA. Our finding that recruitment of Ctf18 to the NPC increased in HU-treated cells (Fig. S2) suggests that the association is important to resolve stalled forks. The unresolved replication fork can lead to fork collapse affecting genomic integrity (Alexander and Orr-Weaver, 2016). Although we found reduced levels of PCNA on stalled replication forks in the *nup170*Δ strain, it remains unclear whether this results from inefficient loading of new PCNA by the Ctf18-RFC complex or if it is a consequence of replication fork collapse. Notwithstanding the underlying reason, reduced PCNA levels on DNA can greatly affect genome stability and chromatin functions.

In *ctf18*Δ cells, the formation of repressor/activator site binding protein 1 (Rap1) foci is altered, which is required to recruit a sirtuin complex - consisting of Sir2, Sir3 and Sir4 - to telomeric DNA for heterochromatin formation and subtelomeric silencing (Hiraga et al., 2006). Similarly, PCNA and CAF-1 have been implicated in heterochromatin establishment and subtelomeric silencing (Kaufman et al., 1997; Miller et al., 2008; Zhang et al., 2000). Mutations in CAF-1 result in a dramatic reduction in DNA-bound histone H3 levels, which leads to reduced levels of Sir2 and Sir4 at silent loci (Tamburini et al., 2006). DNA damage also affects Rap1 and sirtuin-complex association with subtelomeric DNA and triggers loss of subtelomeric silencing (Martin et al., 1999; McAinsh et al., 1999), which could be linked to altered levels of PCNA. We have previously shown that Nup170 physically and functionally interacts with Rap1 and Sir4, and is required for Sir4 assembly on subtelomeric chromatin (Van de Vosse et al., 2013). Taken together, these data point towards a mechanism in which the NPC plays a role in establishing heterochromatin by regulating the Ctf18-RFC complex activities through the regulation of PCNA levels on DNA. A decline in PCNA levels on DNA in the absence of Nup170 can affect the initiation of CAF-1 mediated heterochromatin formation including recruitment of Sir4 to subtelomeric chromatin, which in turn would result in the loss of subtelomeric silencing. This conclusion is further supported by our finding that increasing PCNA levels on DNA by removing Elg1 largely rescues the *nup170*Δ silencing defect.

## MATERIALS and METHODS

### Yeast strains and growth conditions

Unless otherwise specified, all yeast strains used in this study are derived from the S288C background and are listed in Table S2. Cells were grown in YPD medium (1% yeast extract, 2% peptone, and 2% glucose) or synthetic complete (SC) medium containing 2% glucose. To arrest cells in G1, cell cultures were grown to an OD_600_ of 0.6, then α-factor (Sigma-Aldrich, T6901) was added (5g/ml and 100ng/ml for *BAR1* and *bar1*Δ cells respectively) and grown for 2.5 h. Cells were washed twice in YPD and then resuspended in YPD, and samples were harvested at various timepoints. All experiments were performed at 30 °C. Yeast transformations were performed using the lithium acetate/polyethylene glycol method (Daniel Gietz and Woods, 2002). Strains harboring genomic insertions and deletions were made using a plasmid/PCR-based one-step genomic integration method and correct integration was verified by PCR using gene specific primers. The following plasmids were used to construct strains: pJR200 for 3XFLAG tagging (Ranish et al., 2004); pYM14 for 6HA tagging (Janke et al., 2004); pSAN03 for zeocin based deletion (Kumar et al., 2018); and pSJ1256 for GFP_1-10_ tagging (Smoyer et al., 2016). For GFP_11_ tagging, the natR cassette was amplified from pYM42 (Janke et al., 2004) using a forward primer (5′- GGTGGAGGTTCTGGAGGAGGTAGTAGAGATCATATGGTTTTGCATGAATATGTTAATGCTGCTGGTATTACTTAAGGATCCCCGGGTTAATT-3′) containing GFP_11_ and a linker sequence and a reverse primer (5′-AGCTCGATTACAACAGGTGT-3′). The resulting 1393 base pair PCR fragment was used as a template for gene-specific tagging.

### Transcriptome correlation analysis

Using the deleteome compendium of 1,484 yeast deletion transcriptomic profiles (Kemmeren et al., 2014), we identified gene deletion strains with profiles similar to the *nup170*Δ strain. This was done by first listing the genes in the signature of the *nup170*Δ strain (genes with expression values that differed significantly from WT strains) and then performing correlation tests to compare their log_2_ fold-change values against those from each deletion strain in the deleteome. For strains showing a significant positive correlation with the *nup170*Δ strain, we then performed a reciprocal correlation test where the signature of the deletion strain was correlated against the corresponding genes in the *nup170*Δ profile. Deletion strains showing significant correlations in both cases (FDR-corrected *P*-values < 0.05) were considered similar to the *nup170*Δ strain. To help eliminate potential false positives and focus on high-confidence results, we selected strains whose correlation with the *nup170*Δ signature had an FDR-corrected *P*-value in the lowest 5% of all such values. We found that 40 deletion strains passed these criteria, suggesting that the deleted proteins in these strains may have functional overlap with *NUP170* because their deletion resulted in transcriptomic shifts similar to those in the *nup170*Δ strain. All analyses on deleteome strain gene expression data, including all correlation tests, were performed using the R software package.

### Affinity Purification

To perform affinity capture of C-terminally GFP tagged proteins, cells were grown in YPD to a final OD_600_ of 1 - 1.5 and harvested by centrifugation. The cell pellets were washed twice with 20 mM HEPES-KOH, pH 7.4 followed by a wash with 20 mM HEPES-KOH, pH 7.4, 1.2% polyvinylpyrrolidone, 1:100 protease inhibitor cocktail. The cell pellet was pushed through a syringe directly into liquid nitrogen to flash freeze and make “yeast noodles”. Yeast noodles were cryomilled into fine powder using a 50-ml stainless steel jar and a ball mill (Retsch PM100 Planetary Ball Mill, Haan, Germany). To prevent sample heating the milling jar was placed inside a custom-made Teflon jar insulator during the entire milling process. Each sample was subjected to three cycles of milling (6 min each at 450 rpm with reverse rotation every 45 s, with immersion in liquid nitrogen between each cycle) and the resulting yeast powder was stored at -80 °C. Unless specified otherwise, yeast protein lysates were prepared by resuspending 200 mg of cell powder in 450 µl of extraction buffer containing 20 mM HEPES Buffer (pH 7.4), 110 mM CH_3_CO_2_K, 2 mM MgCl_2_, 100 mM NaCl, antifoam-B emulsion (1:5000), 1:100 protease inhibitor cocktail, 1% Triton X-100. When scaling up, the same ratio of cell powder to extraction buffer was maintained. Cell powder was homogenously dispersed by sonication (4 °C, 1 A, 60 s at 10 s intervals) using a QSonica Q700 equipped with a 4-tip microprobe. The resuspension was clarified by centrifugation at 16,000 *g* for 10 min at 4 °C The clarified protein lysate was mixed with 1.5 mg magnetic beads (Dynabead M-270 Epoxy, ThermoFisher) conjugated with in-house GFP nanobody (Fridy et al., 2014), and the mixture was incubated with gentle agitation for 20 min at 4 °C. The beads were washed four times with the extraction buffer. The captured protein complexes were eluted from the beads by adding 35 µl 1.5× LDS sample buffer (NuPAGE) and incubated at room temperature for 10 min. The load samples from the cleared cell lysates were prepared by TCA precipitation.

### Proteomics sample preparation and LC/MS analysis

#### In-gel digestion

Protein complexes from the eluate were resolved by SDS-PAGE and stained with Coomassie blue (Imperial Protein Stain, ThermoFisher Scientific). Protein bands of interest between 250 kDa and 20 kDa were excised, and the gel slices were transferred into 3× washed 0.5 ml microcentrifuge tubes with slits on the bottom. These shredder tubes were then placed into 1.5 ml collector tubes and the assembly was spun in a benchtop centrifuge at 20,000 *g* until all gel slices were shredded and collected into the collector tubes. Gel pieces were de-stained by incubating in 100 µl of 100 mM ammonium bicarbonate/acetonitrile (1:1 vol/vol) for 30 min followed by 5 min incubation in 500 µl of neat acetonitrile, and the supernatant solution was discarded. The destained gel pieces were submerged in 10 mM DTT/100 mM ammonium bicarbonate solution for 30 min at 56 °C on a thermomixer, followed by washing with neat acetonitrile as before to remove the reducing solution. Immediately, freshly prepared 55 mM iodoacetamide / 100 mM ammonium bicarbonate solution was added to the gel pieces to carry out alkylation in the dark for 40 min followed by a wash in neat acetonitrile. Gel pieces were dried in a speed-vac for 20 min and were re-hydrated in 50 µl of sequencing grade trypsin (Promega, 1.3 ng/µl solution in 10 mM ammonium bicarbonate / 10% acetonitrile). After the trypsin solution was completely absorbed, the gel pieces were completely covered in a minimal required volume of 10 mM ammonium bicarbonate / 10% acetonitrile and tryptic digestion was carried out overnight at 37 °C in an air circulating incubator. The next day, a 1:2 (vol/vol) of 5% formic acid in acetonitrile was added to each tube and incubated for 15 min at 37 °C. Supernatants containing extracted peptides were then transferred into fresh tubes, and gel pieces were re-suspended into 100 µl of 1:1 acetonitrile / 5% formic acid for the final extraction, incubated on a shaker for 10 min at 37 °C, spun down quickly and the second supernatants were carefully transferred into the same tubes containing the first extract. The digests were dried completely into a speed-vac for 10 min, re-suspended in 0.1% Trifluoroacetic acid (TFA) and were desalted using C18 Zip tips (Millipore), following the manufacturer’s protocol. The desalted peptide digests were dried in a speed vac and analyzed by LC/MS.

#### LC/MS analysis and protein search

The LC/MS analysis was performed at the Proteomics Core (Fred Hutch, Seattle WA). Peptides were analyzed by LC/ESI MS/MS with a Thermo Scientific Easy-nLC II (Thermo Scientific, Waltham, MA) nano HPLC system coupled to a hybrid Orbitrap Elite ETD (Thermo Scientific, Waltham, MA) mass spectrometer. In-line de-salting was accomplished using a reversed-phase trap column (100 μm × 20 mm) packed with Magic C_18_AQ (5-μm 200Å resin; Michrom Bioresources, Auburn, CA) followed by peptide separations on a reversed-phase column (75 μm × 250 mm) packed with Magic C_18_AQ (5-μm 100Å resin; Michrom Bioresources, Auburn, CA) directly mounted on the electrospray ion source. A 60 min gradient from 7% to 35% acetonitrile in 0.1% formic acid at a flow rate of 400 nL/min was used for chromatographic separations. The heated capillary temperature was set to 300 °C and a spray voltage of 2,750 V was applied to the electrospray tip. The Orbitrap Elite instrument was operated in the data-dependent mode, switching automatically between MS survey scans in the Orbitrap (AGC target value 1,000,000, resolution 120,000, and injection time 250 ms) with MS/MS spectra acquisition in the dual linear ion trap. The 10 most intense ions from the Fourier-transform (FT) full scan were selected for fragmentation in the dual linear ion trap by collision induced dissociation with a normalized collision energy of 35%. Selected ions were dynamically excluded for 10 s with a list size of 500 and exclusion mass by mass width +/- 10 ppm. Raw LC/MS data were searched with MaxQuant (v.1.6.1.0) (Cox and Mann, 2008) against a custom protein database comprised of 6,729 total yeast protein sequences. MaxQuant search parameters were as follows: peptide spectral match (PSM) false discovery rate (FSD) = 0.01; Protein identification FSD = 0.01; fixed peptide modifications = Oxidation (M) and Acetylation (Protein N-terminal); MS/MS tolerance = 0.5 Da. The common coeluting *Saccharomyces cerevisiae* contaminants as listed on the CRAPOME (Mellacheruvu et al., 2013) were excluded from the list and proteins that were identified with 4 or more unique peptides were included in downstream analyses. Nuclear proteins were identified based on their cellular component annotation on PantherDB (Mi et al., 2021).

### Immunoblotting

Proteins were separated on 4-12% gradient SDS-PAGE gels and transferred to nitrocellulose membranes. Membranes were blocked in blocking buffer (Tris-buffered saline with 0.1% Tween (TBST), with 4% milk powder) and the following antibodies were used for immunodetection: mouse monoclonal anti-FLAG-HRP (Sigma, A8592), mouse monoclonal anti-HA (Sigma, H3663), rabbit polyclonal anti-Clb2 (Santa Cruz; sc-9071), and rabbit polyclonal anti-G-6-PDH (Sigma, A9521). When required, an appropriate HRP conjugated secondary antibody was used.

### ChIP, ChIP-qPCR, and ChIP-seq

Chromatin immunoprecipitation was performed as previously described (Smith et al., 2007; Wan et al., 2009) with the following modifications. Briefly, Cells were arrested in G1, washed twice with YPD and resuspended in YPD containing 200 mM hydroxyurea followed by incubation for 40 min at 30 °C. The cells were crosslinked with 1% formaldehyde and the crosslinking reaction was then quenched by incubation with 125 mM glycine. Cells were then disrupted by a BeadBeater using glass beads and then chromatin was sheared to an average size of ∼400 bp using a Covaris S2 sonicator and the cell lysate was clarified by centrifugation at 16,000 *g*. The DNA concentration in each sample was measured using a Qubit Flex Fluorometer (ThermoFisher). Anti-PCNA antibody (GeneTex, GTX64144) was conjugated with Dynabeads Protein G (ThermoFisher,10004D) per the manufacturer’s instructions. The sheared chromatin (2 µg DNA in 400 µl) was incubated with 1.5 g of the conjugated beads overnight on a rotating platform at 4 °C. The eluate and input samples were reverse crosslinked, treated with Proteinase K, and the DNA was prepared using the ChIP DNA Clean & Concentrator kit (Zymo research cat# D5205). The purified DNA samples were used for ChIP-qPCR and ChIP seq analyses. Real-time qPCR was performed using QuantStudio 5 Real-Time PCR System (ThermoFisher) and PowerUp SYBR™ green master mix. Primers for ARS605, ARS606, ARS607 and ARS609 were previously described (Liu et al., 2020). For ChIP-seq, the library was prepared using NEBNext Ultra II DNA Library Prep Kit (Illumina Cat# E7645S) and the sequencing was performed using an Illumina HiSeq 4000. Paired-end reads were aligned to the yeast genome (Ensembl version 103) using STAR(Dobin et al., 2013) and alignment files were filtered with SAMtools (Li et al., 2009) using the 1804 flag and excluding reads with MAPQ scores below 30 as well as orphan reads and read pairs mapping to different chromosomes. Duplicate reads were marked and removed using picard [How to cite: https://github.com/broadinstitute/picard#citing] and samtools. Alignment files for immunoprecipitated and non-immunoprecipitated control samples containing read pairs passing these criteria were then input to macs2 (Zhang et al., 2008) for peak calling and to generate normalized, bedGraph-formatted read pileup profiles for visualization in the Integrative Genomics Viewer (Robinson et al., 2011).

### RNA isolation and RT-qPCR

Cells were harvested from exponentially growing cultures and the RNA was isolated using MasterPure Yeast RNA Purification Kit. The cDNA was prepared using SuperScript III First-Strand Synthesis System (ThermoFisher). The real-time qPCR was performed as described above using gene specific primer pairs listed in Table S3. *RFC1* was used as a reference gene and the data was analyzed using ThermoFisher’s Relative Quantification (RQ) tool.

### Sister Chromatid Cohesion Assay

LacI-GFP/lac operator repeat strains (Mayer et al., 2001; Sanchez et al., 1999) were grown logarithmically in YPD, cells were collected and washed with synthetic complete (SC) media lacking histidine. After washing, cells were resuspended in SC media containing 15 mM 3-amino-1,2,4-triazole (3AT; Sigma) and 5 µg/ml α-factor (Sigma-Aldrich, T6901), and grown for 2.5 h. After G1 arrest, cells were resuspended in YPD containing 15 µg/ml nocodazole (Sigma-Aldrich, M1404) to arrest them in G2/M. Cells were fixed in 4% paraformaldehyde and the number of GFP spots in each cell was scored.

### Fluorescence microscopy and quantification

Images were acquired on a DeltaVision Elite high-resolution microscope (GE Healthcare) equipped with a 100× 1.4 NA objective lens (Olympus). Fluorescence excitation was driven by an Insight SSI solid state light engine (Cytiva) and fluorescence emission was collected by a CoolSnap HQ2 CCD camera (Photometrics). Acquired images were deconvolved using Huygens Professional Software (Scientific Volume Imaging BV, The Netherlands) using theoretically determined point spread functions. For the cell cycle colocalization experiment, cells co-expressing Ctf18-GFP and Nup188-mCherry were grown in YPD and arrested in G1 by adding α factor. After washing the cells, a slide was prepared as described previously (Mast et al., 2016) and images were acquired at regular intervals for 110 min. Object-based colocalization analysis was performed using Imaris (Bitplane). The florescence signal from Nup188-mCherry was processed with the “Surface” function and the Ctf18-GFP fluorescence signal was processed with the “spots” functions. The “Find Spots Close to Surface” function was used to identify Ctf18-GFP spots proximal to Nup188-mCherry.

## Acknowledgement

We thank Philip Hieter and Jeff Ranish for their kind gifts of reagents. We thank Phil Gafken and Lisa Jones of Proteomics & Metabolomics Shared Resource of Fred Hutch for help in MS analysis. We thank Paul Olivier and other members of the Aitchison laboratory for thoughtful discussions. This work was supported by grants P41GM109824 and R01 GM112108 from the National Institutes of Health, to JDA and MPR.

## Notes

### Competing Interest Statement

The authors have declared no competing interest.

## REFERENCES

Aitchison, J.D., and Rout, M.P. (2012). The Yeast Nuclear Pore Complex and Transport Through It. Genetics 190, 855–883. https://doi.org/10.1534/genetics.111.127803.

Akey, C.W., Singh, D., Ouch, C., Echeverria, I., Nudelman, I., Varberg, J.M., Yu, Z., Fang, F., Shi, Y., Wang, J., et al. (2022). Comprehensive structure and functional adaptations of the yeast nuclear pore complex. Cell 185, 361-378.e25. https://doi.org/10.1016/j.cell.2021.12.015.

Alber, F., Dokudovskaya, S., Veenhoff, L.M., Zhang, W., Kipper, J., Devos, D., Suprapto, A., Karni-Schmidt, O., Williams, R., Chait, B.T., et al. (2007). The molecular architecture of the nuclear pore complex. Nature 450, 695–701. https://doi.org/10.1038/nature06405.

Alexander, J.L., and Orr-Weaver, T.L. (2016). Replication fork instability and the consequences of fork collisions from rereplication. Genes Dev. 30, 2241–2252. https://doi.org/10.1101/gad.288142.116.

Alvino, G.M., Collingwood, D., Murphy, J.M., Delrow, J., Brewer, B.J., and Raghuraman, M.K. (2007). Replication in Hydroxyurea: It’s a Matter of Time. Mol Cell Biol 27, 6396–6406. https://doi.org/10.1128/MCB.00719-07.

Baker, R.T., Tobias, J.W., and Varshavsky, A. (1992). Ubiquitin-specific proteases of Saccharomyces cerevisiae. Cloning of UBP2 and UBP3, and functional analysis of the UBP gene family. Journal of Biological Chemistry 267, 23364–23375. https://doi.org/10.1016/S0021-9258(18)50100-9.

Bellaoui, M., Chang, M., Ou, J., Xu, H., Boone, C., and Brown, G.W. (2003). Elg1 forms an alternative RFC complex important for DNA replication and genome integrity. EMBO J 22, 4304–4313. https://doi.org/10.1093/emboj/cdg406.

Bermudez, V.P., Maniwa, Y., Tappin, I., Ozato, K., Yokomori, K., and Hurwitz, J. (2003). The alternative Ctf18-Dcc1-Ctf8-replication factor C complex required for sister chromatid cohesion loads proliferating cell nuclear antigen onto DNA. PNAS 100, 10237–10242. https://doi.org/10.1073/pnas.1434308100.

Boehm, E.M., Gildenberg, M.S., and Washington, M.T. (2016). The many roles of PCNA in eukaryotic DNA replication. Enzymes 39, 231–254. https://doi.org/10.1016/bs.enz.2016.03.003.

Bowman, G.D., O’Donnell, M., and Kuriyan, J. (2004). Structural analysis of a eukaryotic sliding DNA clamp–clamp loader complex. Nature 429, 724–730. https://doi.org/10.1038/nature02585.

Breuer, M., and Ohkura, H. (2015). A negative loop within the nuclear pore complex controls global chromatin organization. Genes Dev. 29, 1789–1794. https://doi.org/10.1101/gad.264341.115.

Brickner, D.G., Randise-Hinchliff, C., Corbin, M.L., Liang, J.M., Kim, S., Sump, B., D’Urso, A., Kim, S.H., Satomura, A., Schmit, H., et al. (2019). The Role of Transcription Factors and Nuclear Pore Proteins in Controlling the Spatial Organization of the Yeast Genome. Developmental Cell 49, 936-947.e4. https://doi.org/10.1016/j.devcel.2019.05.023.

Bylund, G.O., and Burgers, P.M.J. (2005). Replication Protein A-Directed Unloading of PCNA by the Ctf18 Cohesion Establishment Complex. Molecular and Cellular Biology 25, 5445–5455. https://doi.org/10.1128/MCB.25.13.5445-5455.2005.

Cabantous, S., and Waldo, G.S. (2006). In vivo and in vitro protein solubility assays using split GFP. Nature Methods 3, 845–854. https://doi.org/10.1038/nmeth932.

Cox, J., and Mann, M. (2008). MaxQuant enables high peptide identification rates, individualized p.p.b.-range mass accuracies and proteome-wide protein quantification. Nat Biotechnol 26, 1367–1372. https://doi.org/10.1038/nbt.1511.

Crabbé, L., Thomas, A., Pantesco, V., De Vos, J., Pasero, P., and Lengronne, A. (2010). Analysis of replication profiles reveals key role of RFC-Ctf18 in yeast replication stress response. Nat Struct Mol Biol 17, 1391–1397. https://doi.org/10.1038/nsmb.1932.

Daniel Gietz, R., and Woods, R.A. (2002). Transformation of yeast by lithium acetate/single-stranded carrier DNA/polyethylene glycol method. In Methods in Enzymology, C. Guthrie, and G.R. Fink, eds. (Academic Press), pp. 87–96.

Dilworth, D.J., Suprapto, A., Padovan, J.C., Chait, B.T., Wozniak, R.W., Rout, M.P., and Aitchison, J.D. (2001). Nup2p Dynamically Associates with the Distal Regions of the Yeast Nuclear Pore Complex. Journal of Cell Biology 153, 1465–1478. https://doi.org/10.1083/jcb.153.7.1465.

Dobin, A., Davis, C.A., Schlesinger, F., Drenkow, J., Zaleski, C., Jha, S., Batut, P., Chaisson, M., and Gingeras, T.R. (2013). STAR: ultrafast universal RNA-seq aligner. Bioinformatics 29, 15–21. https://doi.org/10.1093/bioinformatics/bts635.

Essers, J., Theil, A.F., Baldeyron, C., van Cappellen, W.A., Houtsmuller, A.B., Kanaar, R., and Vermeulen, W. (2005). Nuclear Dynamics of PCNA in DNA Replication and Repair. Molecular and Cellular Biology 25, 9350–9359. https://doi.org/10.1128/MCB.25.21.9350-9359.2005.

Fernandez-Martinez, J., and Rout, M.P. (2021). One Ring to Rule them All? Structural and Functional Diversity in the Nuclear Pore Complex. Trends in Biochemical Sciences 0. https://doi.org/10.1016/j.tibs.2021.01.003.

Fridy, P.C., Li, Y., Keegan, S., Thompson, M.K., Nudelman, I., Scheid, J.F., Oeffinger, M., Nussenzweig, M.C., Fenyö, D., Chait, B.T., et al. (2014). A robust pipeline for rapid production of versatile nanobody repertoires. Nature Methods 11, 1253–1260. https://doi.org/10.1038/nmeth.3170.

Fujisawa, R., Ohashi, E., Hirota, K., and Tsurimoto, T. (2017). Human CTF18-RFC clamp-loader complexed with non-synthesising DNA polymerase ε efficiently loads the PCNA sliding clamp. Nucleic Acids Research 45, 4550–4563. https://doi.org/10.1093/nar/gkx096.

Gaillard, P.-H.L., Martini, E.M.-D., Kaufman, P.D., Stillman, B., Moustacchi, E., and Almouzni, G. (1996). Chromatin Assembly Coupled to DNA Repair: A New Role for Chromatin Assembly Factor I. Cell 86, 887–896. https://doi.org/10.1016/S0092-8674(00)80164-6.

Galy, V., Olivo-Marin, J.-C., Scherthan, H., Doye, V., Rascalou, N., and Nehrbass, U. (2000). Nuclear pore complexes in the organization of silent telomeric chromatin. Nature 403, 108–112. https://doi.org/10.1038/47528.

Galy, V., Gadal, O., Fromont-Racine, M., Romano, A., Jacquier, A., and Nehrbass, U. (2004). Nuclear Retention of Unspliced mRNAs in Yeast Is Mediated by Perinuclear Mlp1. Cell 116, 63–73. https://doi.org/10.1016/S0092-8674(03)01026-2.

Gellon, L., Razidlo, D.F., Gleeson, O., Verra, L., Schulz, D., Lahue, R.S., and Freudenreich, C.H. (2011). New Functions of Ctf18-RFC in Preserving Genome Stability outside Its Role in Sister Chromatid Cohesion. PLOS Genetics 7, e1001298. https://doi.org/10.1371/journal.pgen.1001298.

Grabarczyk, D.B., Silkenat, S., and Kisker, C. (2018). Structural Basis for the Recruitment of Ctf18-RFC to the Replisome. Structure 26, 137-144.e3. https://doi.org/10.1016/j.str.2017.11.004.

Hakhverdyan, Z., Domanski, M., Hough, L.E., Oroskar, A.A., Oroskar, A.R., Keegan, S., Dilworth, D.J., Molloy, K.R., Sherman, V., Aitchison, J.D., et al. (2015). Rapid, optimized interactomic screening. Nat Methods 12, 553–560. https://doi.org/10.1038/nmeth.3395.

Hanna, J.S., Kroll, E.S., Lundblad, V., and Spencer, F.A. (2001). Saccharomyces cerevisiae CTF18 and CTF4 Are Required for Sister Chromatid Cohesion. Molecular and Cellular Biology 21, 3144–3158. https://doi.org/10.1128/MCB.21.9.3144-3158.2001.

Hiraga, S., Robertson, E.D., and Donaldson, A.D. (2006). The Ctf18 RFC-like complex positions yeast telomeres but does not specify their replication time. The EMBO Journal 25, 1505–1514. https://doi.org/10.1038/sj.emboj.7601038.

Ho, B., Baryshnikova, A., and Brown, G.W. (2018). Unification of Protein Abundance Datasets Yields a Quantitative Saccharomyces cerevisiae Proteome. Cels 6, 192-205.e3. https://doi.org/10.1016/j.cels.2017.12.004.

Hu, C.-D., and Kerppola, T.K. (2003). Simultaneous visualization of multiple protein interactions in living cells using multicolor fluorescence complementation analysis. Nat Biotechnol 21, 539–545. https://doi.org/10.1038/nbt816.

Hu, C.-D., Chinenov, Y., and Kerppola, T.K. (2002). Visualization of Interactions among bZIP and Rel Family Proteins in Living Cells Using Bimolecular Fluorescence Complementation. Molecular Cell 9, 789–798. https://doi.org/10.1016/S1097-2765(02)00496-3.

Janke, C., Magiera, M.M., Rathfelder, N., Taxis, C., Reber, S., Maekawa, H., Moreno-Borchart, A., Doenges, G., Schwob, E., Schiebel, E., et al. (2004). A versatile toolbox for PCR-based tagging of yeast genes: new fluorescent proteins, more markers and promoter substitution cassettes. Yeast 21, 947–962. https://doi.org/10.1002/yea.1142.

Kadota, S., Ou, J., Shi, Y., Lee, J.T., Sun, J., and Yildirim, E. (2020). Nucleoporin 153 links nuclear pore complex to chromatin architecture by mediating CTCF and cohesin binding. Nat Commun 11, 2606. https://doi.org/10.1038/s41467-020-16394-3.

Kaufman, P.D., Kobayashi, R., and Stillman, B. (1997). Ultraviolet radiation sensitivity and reduction of telomeric silencing in Saccharomyces cerevisiae cells lacking chromatin assembly factor-I. Genes Dev. 11, 345–357. https://doi.org/10.1101/gad.11.3.345.

Kehat, I., Accornero, F., Aronow, B.J., and Molkentin, J.D. (2011). Modulation of chromatin position and gene expression by HDAC4 interaction with nucleoporins. J Cell Biol 193, 21–29. https://doi.org/10.1083/jcb.201101046.

Kemmeren, P., Sameith, K., van de Pasch, L.A.L., Benschop, J.J., Lenstra, T.L., Margaritis, T., O’Duibhir, E., Apweiler, E., van Wageningen, S., Ko, C.W., et al. (2014). Large-Scale Genetic Perturbations Reveal Regulatory Networks and an Abundance of Gene-Specific Repressors. Cell 157, 740–752. https://doi.org/10.1016/j.cell.2014.02.054.

Kerscher, O., Hieter, P., Winey, M., and Basrai, M.A. (2001). Novel Role for a Saccharomyces cerevisiae Nucleoporin, Nup170p, in Chromosome Segregation. Genetics 157, 1543–1553..

Kim, S.J., Fernandez-Martinez, J., Nudelman, I., Shi, Y., Zhang, W., Raveh, B., Herricks, T., Slaughter, B.D., Hogan, J.A., Upla, P., et al. (2018). Integrative structure and functional anatomy of a nuclear pore complex. Nature 555, 475–482. https://doi.org/10.1038/nature26003.

Kubota, T., Hiraga, S., Yamada, K., Lamond, A.I., and Donaldson, A.D. (2011). Quantitative Proteomic Analysis of Chromatin Reveals that Ctf18 Acts in the DNA Replication Checkpoint. Mol Cell Proteomics 10, M110.005561. https://doi.org/10.1074/mcp.M110.005561.

Kubota, T., Nishimura, K., Kanemaki, M.T., and Donaldson, A.D. (2013). The Elg1 Replication Factor C-like Complex Functions in PCNA Unloading during DNA Replication. Molecular Cell 50, 273–280. https://doi.org/10.1016/j.molcel.2013.02.012.

Kumar, S., de Boer, R., and van der Klei, I.J. (2018). Yeast cells contain a heterogeneous population of peroxisomes that segregate asymmetrically during cell division. Journal of Cell Science 131, jcs207522. https://doi.org/10.1242/jcs.207522.

Lamm, N., Rogers, S., and Cesare, A.J. (2021). Chromatin mobility and relocation in DNA repair. Trends in Cell Biology 31, 843–855. https://doi.org/10.1016/j.tcb.2021.06.002.

Lapetina, D.L., Ptak, C., Roesner, U.K., and Wozniak, R.W. (2017). Yeast silencing factor Sir4 and a subset of nucleoporins form a complex distinct from nuclear pore complexes. J Cell Biol 216, 3145–3159. https://doi.org/10.1083/jcb.201609049.

Lengronne, A., McIntyre, J., Katou, Y., Kanoh, Y., Hopfner, K.-P., Shirahige, K., and Uhlmann, F. (2006). Establishment of Sister Chromatid Cohesion at the S. cerevisiae Replication Fork. Molecular Cell 23, 787–799. https://doi.org/10.1016/j.molcel.2006.08.018.

Li, H., Handsaker, B., Wysoker, A., Fennell, T., Ruan, J., Homer, N., Marth, G., Abecasis, G., Durbin, R., and 1000 Genome Project Data Processing Subgroup (2009). The Sequence Alignment/Map format and SAMtools. Bioinformatics 25, 2078–2079. https://doi.org/10.1093/bioinformatics/btp352.

Light, W.H., Brickner, D.G., Brand, V.R., and Brickner, J.H. (2010). Interaction of a DNA Zip Code with the Nuclear Pore Complex Promotes H2A.Z Incorporation and INO1 Transcriptional Memory. Molecular Cell 40, 112–125. https://doi.org/10.1016/j.molcel.2010.09.007.

Liu, H.W., Bouchoux, C., Panarotto, M., Kakui, Y., Patel, H., and Uhlmann, F. (2020). Division of Labor between PCNA Loaders in DNA Replication and Sister Chromatid Cohesion Establishment. Molecular Cell 78, 725-738.e4. https://doi.org/10.1016/j.molcel.2020.03.017.

Mailand, N., Gibbs-Seymour, I., and Bekker-Jensen, S. (2013). Regulation of PCNA–protein interactions for genome stability. Nat Rev Mol Cell Biol 14, 269–282. https://doi.org/10.1038/nrm3562.

Mast, F.D., Jamakhandi, A., Saleem, R.A., Dilworth, D.J., Rogers, R.S., Rachubinski, R.A., and Aitchison, J.D. (2016). Peroxins Pex30 and Pex29 Dynamically Associate with Reticulons to Regulate Peroxisome Biogenesis from the Endoplasmic Reticulum. J Biol Chem 291, 15408–15427. https://doi.org/10.1074/jbc.M116.728154.

Mayer, M.L., Gygi, S.P., Aebersold, R., and Hieter, P. (2001). Identification of RFC(Ctf18p, Ctf8p, Dcc1p): An Alternative RFC Complex Required for Sister Chromatid Cohesion in S. cerevisiae. Molecular Cell 7, 959–970. https://doi.org/10.1016/S1097-2765(01)00254-4.

Mellacheruvu, D., Wright, Z., Couzens, A.L., Lambert, J.-P., St-Denis, N.A., Li, T., Miteva, Y.V., Hauri, S., Sardiu, M.E., Low, T.Y., et al. (2013). The CRAPome: a contaminant repository for affinity purification–mass spectrometry data. Nature Methods 10, 730–736. https://doi.org/10.1038/nmeth.2557.

Mi, H., Ebert, D., Muruganujan, A., Mills, C., Albou, L.-P., Mushayamaha, T., and Thomas, P.D. (2021). PANTHER version 16: a revised family classification, tree-based classification tool, enhancer regions and extensive API. Nucleic Acids Research 49, D394–D403. https://doi.org/10.1093/nar/gkaa1106.

Miller, A., Yang, B., Foster, T., and Kirchmaier, A.L. (2008). Proliferating Cell Nuclear Antigen and ASF1 Modulate Silent Chromatin in Saccharomyces cerevisiae via Lysine 56 on Histone H3. Genetics 179, 793–809. https://doi.org/10.1534/genetics.107.084525.

Moggs, J.G., Grandi, P., Quivy, J.-P., Jónsson, Z.O., Hübscher, U., Becker, P.B., and Almouzni, G. (2000). A CAF-1–PCNA-Mediated Chromatin Assembly Pathway Triggered by Sensing DNA Damage. Molecular and Cellular Biology 20, 1206–1218. https://doi.org/10.1128/MCB.20.4.1206-1218.2000.

Naiki, T., Kondo, T., Nakada, D., Matsumoto, K., and Sugimoto, K. (2001). Chl12 (Ctf18) Forms a Novel Replication Factor C-Related Complex and Functions Redundantly with Rad24 in the DNA Replication Checkpoint Pathway. Molecular and Cellular Biology 21, 5838–5845. https://doi.org/10.1128/MCB.21.17.5838-5845.2001.

Niepel, M., Strambio-de-Castillia, C., Fasolo, J., Chait, B.T., and Rout, M.P. (2005). The nuclear pore complex–associated protein, Mlp2p, binds to the yeast spindle pole body and promotes its efficient assembly. Journal of Cell Biology 170, 225–235. https://doi.org/10.1083/jcb.200504140.

Ogiwara, H., Ohuchi, T., Ui, A., Tada, S., Enomoto, T., and Seki, M. (2007). Ctf18 is required for homologous recombination-mediated double-strand break repair. Nucleic Acids Res 35, 4989–5000. https://doi.org/10.1093/nar/gkm523.

Ptak, C., Aitchison, J.D., and Wozniak, R.W. (2014). The multifunctional nuclear pore complex: a platform for controlling gene expression. Curr Opin Cell Biol 0, 46–53. https://doi.org/10.1016/j.ceb.2014.02.001.

Ranish, J.A., Hahn, S., Lu, Y., Yi, E.C., Li, X., Eng, J., and Aebersold, R. (2004). Identification of TFB5, a new component of general transcription and DNA repair factor IIH. Nat Genet 36, 707–713. https://doi.org/10.1038/ng1385.

Robinson, J.T., Thorvaldsdóttir, H., Winckler, W., Guttman, M., Lander, E.S., Getz, G., and Mesirov, J.P. (2011). Integrative genomics viewer. Nat Biotechnol 29, 24–26. https://doi.org/10.1038/nbt.1754.

Sanchez, Y., Bachant, J., Wang, H., Hu, F., Liu, D., Tetzlaff, M., and Elledge, S.J. (1999). Control of the DNA Damage Checkpoint by Chk1 and Rad53 Protein Kinases Through Distinct Mechanisms. Science 286, 1166–1171. https://doi.org/10.1126/science.286.5442.1166.

Serra-Cardona, A., and Zhang, Z. (2018). Replication-coupled nucleosome assembly as a passage of epigenetic information and cell identity. Trends Biochem Sci 43, 136–148. https://doi.org/10.1016/j.tibs.2017.12.003.

Smith, J.J., Ramsey, S.A., Marelli, M., Marzolf, B., Hwang, D., Saleem, R.A., Rachubinski, R.A., and Aitchison, J.D. (2007). Transcriptional responses to fatty acid are coordinated by combinatorial control. Mol Syst Biol 3, 115. https://doi.org/10.1038/msb4100157.

Smolikov, S., Mazor, Y., and Krauskopf, A. (2004). ELG1, a regulator of genome stability, has a role in telomere length regulation and in silencing. PNAS 101, 1656–1661. https://doi.org/10.1073/pnas.0307796100.

Smoyer, C.J., Katta, S.S., Gardner, J.M., Stoltz, L., McCroskey, S., Bradford, W.D., McClain, M., Smith, S.E., Slaughter, B.D., Unruh, J.R., et al. (2016). Analysis of membrane proteins localizing to the inner nuclear envelope in living cells. J Cell Biol 215, 575–590. https://doi.org/10.1083/jcb.201607043.

Stark, C., Breitkreutz, B.-J., Reguly, T., Boucher, L., Breitkreutz, A., and Tyers, M. (2006). BioGRID: a general repository for interaction datasets. Nucleic Acids Research 34, D535–D539. https://doi.org/10.1093/nar/gkj109.

Stokes, K., Winczura, A., Song, B., De Piccoli, G., and Grabarczyk, D.B. (2020). Ctf18-RFC and DNA Pol LJ form a stable leading strand polymerase/clamp loader complex required for normal and perturbed DNA replication. Nucleic Acids Research 48, 8128–8145. https://doi.org/10.1093/nar/gkaa541.

Strambio-de-Castillia, C., Blobel, G., and Rout, M.P. (1999). Proteins Connecting the Nuclear Pore Complex with the Nuclear Interior. Journal of Cell Biology 144, 839–855. https://doi.org/10.1083/jcb.144.5.839.

Suter, B., Tong, A., Chang, M., Yu, L., Brown, G.W., Boone, C., and Rine, J. (2004). The Origin Recognition Complex Links Replication, Sister Chromatid Cohesion and Transcriptional Silencing in Saccharomyces cerevisiae. Genetics 167, 579–591. https://doi.org/10.1534/genetics.103.024851.

Tamburini, B.A., Carson, J.J., Linger, J.G., and Tyler, J.K. (2006). Dominant Mutants of the Saccharomyces cerevisiae ASF1 Histone Chaperone Bypass the Need for CAF-1 in Transcriptional Silencing by Altering Histone and Sir Protein Recruitment. Genetics 173, 599–610. https://doi.org/10.1534/genetics.105.054783.

Tan-Wong, S.M., Wijayatilake, H.D., and Proudfoot, N.J. (2009). Gene loops function to maintain transcriptional memory through interaction with the nuclear pore complex. Genes Dev. 23, 2610–2624. https://doi.org/10.1101/gad.1823209.

Van de Vosse, D.W., Wan, Y., Lapetina, D.L., Chen, W.-M., Chiang, J.-H., Aitchison, J.D., and Wozniak, R.W. (2013). A Role for the Nucleoporin Nup170p in Chromatin Structure and Gene Silencing. Cell 152, 969–983. https://doi.org/10.1016/j.cell.2013.01.049.

Wan, Y., Saleem, R.A., Ratushny, A.V., Roda, O., Smith, J.J., Lin, C.-H., Chiang, J.-H., and Aitchison, J.D. (2009). Role of the Histone Variant H2A.Z/Htz1p in TBP Recruitment, Chromatin Dynamics, and Regulated Expression of Oleate-Responsive Genes. Molecular and Cellular Biology 29, 2346–2358. https://doi.org/10.1128/MCB.01233-08.

Wente, S.R., Rout, M.P., and Blobel, G. (1992). A new family of yeast nuclear pore complex proteins. Journal of Cell Biology 119, 705–723. https://doi.org/10.1083/jcb.119.4.705.

Whalen, J.M., and Freudenreich, C.H. (2020). Location, Location, Location: The Role of Nuclear Positioning in the Repair of Collapsed Forks and Protection of Genome Stability. Genes 11, 635. https://doi.org/10.3390/genes11060635.

Yao, N.Y., and O’Donnell, M. (2012). The RFC Clamp Loader: Structure and Function. Subcell Biochem 62, 259–279. https://doi.org/10.1007/978-94-007-4572-8_14.

Young, T.J., Cui, Y., Pfeffer, C., Hobbs, E., Liu, W., Irudayaraj, J., and Kirchmaier, A.L. (2020). CAF-1 and Rtt101p function within the replication-coupled chromatin assembly network to promote H4 K16ac, preventing ectopic silencing. PLOS Genetics 16, e1009226. https://doi.org/10.1371/journal.pgen.1009226.

Zhang, Y., Liu, T., Meyer, C.A., Eeckhoute, J., Johnson, D.S., Bernstein, B.E., Nusbaum, C., Myers, R.M., Brown, M., Li, W., et al. (2008). Model-based analysis of ChIP-Seq (MACS). Genome Biol 9, R137. https://doi.org/10.1186/gb-2008-9-9-r137.

Zhang, Z., Shibahara, K., and Stillman, B. (2000). PCNA connects DNA replication to epigenetic inheritance in yeast. Nature 408, 221–225. https://doi.org/10.1038/35041601.

