## Supplementary material for "The nucleoporin Nup170 mediates subtelomeric gene silencing through the Ctf18-RFC complex and PCNA": Table S1

**Candidates identified by Nup170-GFP AP-MS:**

| Nup170 PPI Uniprot IDs | Nuclear protein without Nups |
| --- | --- |
| P32600 | Y |
| P38811 | Y |
| P40064 | N |
| Q06179 | N |
| P07245 | Y |
| P04801 | N |
| P35169 | Y |
| P35729 | N |
| P31539 | Y |
| P05453 | N |
| P14907 | N |
| P00958 | N |
| P37291 | N |
| P07244 | N |
| P15424 | Y |
| P46655 | N |
| P30624 | N |
| P09440 | N |
| Q03280 | Y |
| Q00402 | N |
| P40825 | N |
| Q05911 | N |
| P19414 | N |
| P40991 | Y |
| P00931 | Y |
| P32861 | N |
| P38088 | N |
| P16603 | N |
| Q02892 | Y |
| P23254 | N |
| P22137 | N |
| P15303 | N |
| P52910 | Y |
| P32563 | N |
| P47075 | N |
| P04806 | N |
| Q12680 | N |
| P07284 | N |
| P06738 | N |
| P43535 | N |
| P31688 | N |
| P39685 | N |
| P40217 | N |
| P27616 | Y |
| P34077 | N |
| P06168 | N |
| P36120 | Y |
| P40469 | Y |
| P47117 | N |
| P53297 | Y |
| P31116 | Y |
| P53742 | Y |
| P07283 | N |
| P14906 | Y |
| P19158 | N |
| P49095 | N |
| P52593 | N |
| Q99207 | Y |
| Q04119 | Y |
| P46673 | N |
| Q03790 | N |
| P53090 | N |
| Q04458 | N |
| P36114 | Y |
| Q07807 | N |
| P18963 | N |
| P37292 | N |
| P09624 | N |
| P49723 | Y |
| Q08001 | N |
| Q12363 | Y |
| Q04177 | Y |
| P38011 | N |
| P25294 | Y |
| P35191 | N |
| P25379 | N |
| P00498 | N |
| P17709 | N |
| Q02725 | N |
| Q04409 | N |
| P07257 | N |
| P21147 | N |
| P35844 | N |
| P37297 | N |
| P39109 | N |
| P40368 | N |
| P54839 | Y |
| Q00416 | Y |
| Q04175 | Y |
| P32794 | N |
| P38625 | N |
| P01120 | Y |
| P41807 | N |
| P47176 | Y |
| P00817 | Y |
| P00942 | N |
| P14020 | N |
| P32356 | N |
| P32767 | Y |
| P40071 | N |
| P46995 | Y |
| Q08951 | N |
| P41805 | N |
| P53941 | Y |
| P48234 | Y |
| P09064 | N |
| Q08096 | Y |
| P38828 | Y |
| P32476 | N |
| P38077 | N |
| P46367 | N |
| P38630 | Y |
| P52489 | N |
| P07213 | N |
| P08524 | N |
| P10614 | N |
| P12868 | N |
| P13663 | Y |
| P25342 | N |
| P32366 | N |
| P36148 | N |
| P38153 | N |
| P38251 | Y |
| P43616 | N |
| P49687 | N |
| P53235 | N |
| Q12074 | Y |
| Q12265 | N |
| P25555 | Y |
| Q12159 | Y |
| P40362 | Y |
| P33201 | Y |
| Q06132 | Y |
| P12754 | N |
| P22224 | N |
| P25623 | N |
| P33307 | Y |
| P33767 | Y |
| P40215 | N |
| Q06205 | Y |
| P22213 | N |
| P25375 | N |
| P32336 | Y |
| P40986 | N |
| P43633 | Y |
| P49956 | Y |
| Q00764 | N |
| Q03653 | N |
| Q04894 | N |
| Q06488 | Y |
| Q12452 | N |
| Q12746 | N |
| P10080 | Y |
| P30822 | Y |
| P19454 | Y |
| P21576 | N |
| P23641 | N |
| P38629 | Y |
| P20606 | N |
| P20449 | Y |
| P38264 | N |
| P23638 | Y |
| P32473 | N |
| P38115 | N |
| P40016 | N |
| Q12154 | N |
| Q12447 | N |
| P38286 | N |
| P16550 | Y |
| P38086 | Y |
| P38810 | N |
| P26785 | N |
| P12688 | N |
| P07246 | N |
| Q02785 | N |
| P05374 | N |
| P12709 | N |
| P32419 | N |
| P33333 | N |
| P40047 | N |
| P40462 | N |
| P40477 | N |
| P40483 | N |
| P41318 | Y |
| P46951 | Y |
| P46956 | N |
| P47120 | Y |
| P53199 | N |
| P53731 | N |
| P54885 | Y |
| Q02046 | Y |
| Q02199 | N |
| Q02805 | N |
| Q05812 | N |
| Q06440 | N |
| Q07551 | Y |
| Q08921 | N |
| P25567 | N |
| P53336 | Y |
| P53145 | N |
| P09436 | N |
| P25087 | N |
| P40348 | Y |
| Q04947 | N |
| P25293 | Y |
| P36165 | N |
| Q05123 | Y |
| Q12028 | N |
| P37263 | Y |
| P32598 | Y |
| P12695 | N |
| P38626 | Y |
| P50085 | N |
| P89102 | N |
| Q05016 | Y |
| P12684 | Y |
| P32466 | N |
| P39002 | N |
| Q06108 | N |
| P08417 | N |
| P08539 | N |
| P20967 | N |
| P25297 | N |
| P27351 | N |
| P32468 | N |
| P32618 | N |
| P38689 | N |
| P40302 | Y |
| P43123 | Y |
| P47025 | N |
| P53852 | N |
| P54783 | N |
| Q02629 | N |
| Q02932 | Y |
| Q06668 | N |
| Q12109 | N |
| Q12246 | N |
| P40495 | N |
| P53335 | Y |
| P40075 | Y |
| Q07362 | Y |
| P39078 | N |
| P47006 | Y |
| P39976 | N |
| Q04067 | N |
| P17505 | N |
| P20459 | N |
| P40961 | N |
| P53327 | Y |
| Q01477 | N |
| P15891 | N |
| P20795 | N |
| P21243 | Y |
| P38174 | Y |
| Q06685 | N |
| Q12114 | N |
| P26784 | N |
| P38891 | N |
| P32386 | N |
| P00815 | N |
| P10869 | N |
| P21560 | N |
| P23180 | N |
| P25349 | N |
| P27514 | N |
| P32775 | N |
| P32833 | Y |
| P34756 | N |
| P38127 | N |
| P38234 | N |
| P38861 | Y |
| P39925 | N |
| P39958 | N |
| P40066 | N |
| P40339 | Y |
| P40350 | N |
| P40468 | N |
| P40547 | Y |
| P41921 | Y |
| P46151 | N |
| P47042 | N |
| Q00245 | N |
| Q03177 | N |
| Q03281 | Y |
| Q04371 | Y |
| Q06010 | N |
| Q08220 | N |
| Q10740 | Y |
| P21242 | Y |
| P32899 | Y |
| P15180 | N |
| Q06287 | Y |
| Q08421 | Y |
| P51601 | Y |
| P15705 | N |
| P26637 | N |
| P39954 | N |
| P15992 | Y |
| P38968 | N |
| P00937 | N |
| P06780 | N |
| P15703 | N |
| P15790 | Y |
| P17423 | N |
| P25605 | N |
| P32614 | N |
| P32915 | N |
| P38065 | N |
| P39007 | N |
| P40531 | N |
| P42842 | Y |
| P46982 | N |
| Q04013 | N |
| Q12018 | Y |
| Q12150 | N |
| P00546 | Y |
| P23337 | N |
| Q03161 | Y |
| P07560 | N |
| P01123 | N |
| P04046 | N |
| P08466 | Y |
| P14743 | N |
| P18562 | N |
| P19263 | Y |
| P19881 | Y |
| P25340 | N |
| P25374 | Y |
| P28321 | N |
| P32288 | Y |
| P32500 | N |
| P36007 | N |
| P36060 | N |
| P38840 | Y |
| P38883 | Y |
| P38972 | N |
| P38985 | Y |
| P38988 | N |
| P40098 | N |
| P40513 | N |
| P40557 | N |
| P42935 | Y |
| P43613 | N |
| P43639 | Y |
| P48445 | Y |
| P52891 | N |
| P53049 | N |
| P53253 | N |
| Q01532 | N |
| Q02724 | Y |
| Q02776 | N |
| Q03441 | N |
| Q04472 | N |
| Q05050 | N |
| Q06508 | N |
| Q08548 | N |
| Q08686 | N |
| Q12445 | N |
| Q99186 | N |
| P43588 | Y |
| P07260 | Y |
| P38805 | Y |
| P02406 | Y |
| P04840 | N |
| P35178 | Y |
| P38712 | Y |
| P21375 | N |
| P32787 | N |
| P53551 | Y |
| P38715 | Y |
| P40509 | N |
| P53598 | N |
| P54860 | Y |
| Q04178 | Y |
| Q08732 | Y |
| Q08959 | N |
| Q12335 | N |
| P32590 | N |
| P05739 | N |
| P26570 | Y |
| P36018 | N |
| P38755 | N |
| P01119 | Y |
| P04076 | N |
| P06197 | N |
| P10662 | N |
| P15274 | N |
| P15873 | Y |
| P17442 | Y |
| P17649 | N |
| P20051 | Y |
| P23639 | Y |
| P27929 | N |
| P30656 | Y |
| P30657 | Y |
| P32331 | N |
| P32458 | N |
| P32901 | N |
| P32939 | N |
| P33775 | N |
| P36010 | N |
| P36022 | N |
| P36037 | Y |
| P36164 | N |
| P38177 | N |
| P38737 | Y |
| P38821 | N |
| P39986 | N |
| P40008 | N |
| P40091 | N |
| P40093 | Y |
| P43590 | N |
| P43620 | N |
| P43636 | N |
| P47050 | Y |
| P47140 | N |
| P48837 | N |
| P50107 | Y |
| P53248 | N |
| P53736 | N |
| P53865 | N |
| P53874 | Y |
| P53981 | Y |
| P54784 | Y |
| Q02206 | Y |
| Q02950 | N |
| Q03516 | N |
| Q05029 | N |
| Q05905 | Y |
| Q06678 | N |
| Q08226 | N |
| Q12072 | Y |
| Q12123 | Y |
| Q12404 | N |
| Q12466 | N |
| Q12514 | Y |
| Q99190 | N |
| P06105 | Y |
| P39730 | N |
| P34247 | Y |
| Q04867 | Y |
| P38431 | N |
| P20434 | Y |
| Q03690 | N |
| P14741 | N |
| P32379 | Y |
| P53137 | N |
| Q08960 | N |
| Q99216 | Y |
| P54837 | N |
| Q12453 | Y |
| P12904 | Y |
| Q08601 | Y |
| Q12125 | N |
| P13517 | N |
| P25334 | N |
| P25368 | Y |
| P32353 | N |
| P32561 | Y |
| P35723 | N |
| P36101 | N |
| P40485 | N |
| P51534 | N |
| P53912 | Y |
| P40510 | N |
| Q02326 | N |
| P12683 | Y |
| P16892 | Y |
| P23585 | N |
| P43585 | N |
| P03965 | N |
| P05373 | Y |
| P07267 | N |
| P10834 | N |
| P11914 | N |
| P11927 | N |
| P13259 | Y |
| P15700 | Y |
| P16622 | N |
| P17558 | N |
| P22141 | Y |
| P25039 | N |
| P25847 | Y |
| P27636 | N |
| P28495 | N |
| P28496 | Y |
| P28817 | N |
| P32179 | Y |
| P32259 | Y |
| P32332 | N |
| P32342 | Y |
| P32566 | Y |
| P32832 | Y |
| P33334 | Y |
| P34248 | N |
| P35193 | N |
| P36528 | N |
| P38226 | N |
| P38230 | Y |
| P38295 | N |
| P38355 | N |
| P38833 | Y |
| P39102 | Y |
| P39714 | N |
| P40363 | N |
| P40582 | N |
| P47818 | N |
| P48353 | N |
| P49435 | Y |
| P50874 | Y |
| P51998 | N |
| P53154 | N |
| P53290 | Y |
| P53292 | N |
| P53866 | Y |
| P53954 | N |
| P53969 | N |
| P80428 | Y |
| Q03002 | Y |
| Q03305 | Y |
| Q03308 | N |
| Q03774 | Y |
| Q04651 | N |
| Q04958 | N |
| Q05359 | N |
| Q05567 | N |
| Q06151 | Y |
| Q06505 | Y |
| Q08985 | Y |
| Q12071 | N |
| Q12096 | N |
| Q12102 | Y |
| Q12133 | N |
| Q12179 | Y |
| Q12207 | N |
| Q12387 | N |
| Q12458 | Y |
| Q99189 | Y |
| Q99394 | N |
| P23776 | N |
| P39013 | N |
| P40059 | Y |
| P38061 | N |
| P28000 | Y |
| Q02979 | N |
| P32368 | N |
| Q12522 | Y |
| Q12339 | Y |
| Q08723 | N |
| Q03640 | N |
| P03872 | N |
| P25582 | Y |
| P53136 | Y |
| P40470 | Y |
| Q12052 | Y |
| Q06511 | Y |
| P12611 | N |
| P23292 | Y |
| P29547 | Y |
| P30619 | N |
| P36057 | N |
| P36144 | Y |
| P36161 | N |
| P38353 | N |
| P38427 | N |
| P41819 | Y |
| P48813 | N |
| P53163 | N |
| P53337 | N |
| Q02555 | Y |
| Q02959 | Y |
| Q03048 | Y |
| Q03088 | N |
| Q05506 | N |
| Q12306 | Y |
