## Supplementary material for "The nucleoporin Nup170 mediates subtelomeric gene silencing through the Ctf18-RFC complex and PCNA": Table S2

**Table S1. Yeast strains**

| **Strain** | **Genotype** | **Reference** |
| --- | --- | --- |
| *BY4741* | *MAT a his3∆1 leu2∆0 ura3∆0 met15∆0* | (Brachmann et al., 1998) |
| *BY4742* | *MATα his3∆1 leu2∆0 ura3∆0 lys2∆0* | (Brachmann et al., 1998) |
| SKY137 | *MATα his3∆1 leu2∆0 ura3∆0 lys2∆0 NUP170-GFP::LEU2* | This study |
| Y819 | *MATa trp1-1:lacO-TRP1-LEU2 his3-11,15:lacR::GFP-HIS3 leu2-3,112 ura3-1 ade2-1 can1-100* | (Mayer et al., 2001) |
| SKY360 | *MATa trp1-1:lacO-TRP1-LEU2 his3-11,15:lacR::GFP-HIS3 leu2-3,112 ura3-1 ade2-1 can1-100 nup170::hph* | Derived from Y819 |
| SKY411 | *MATa trp1-1:lacO-TRP1-LEU2 his3-11,15:lacR::GFP-HIS3 leu2-3,112 ura3-1 ade2-1 can1-100 ctf18::kanR* | Derived from Y819 |
| SKY415 | *Mat a his3Δ1 leu2Δ0 ura3Δ0 met15∆0 CTF18-GFP::HisMX bar1::natR* | This study |
| SKY416 | *Mat a his3Δ1 leu2Δ0 ura3Δ0 met15∆0 CTF18-GFP::HisMX nup170::kanMX bar1::natR* | This study |
| SKY422 | *Mat a his3Δ1 leu2Δ0 ura3Δ0 met15∆0 CTF18-GFP::HisMX bar1::natR NUP170-3FLAG::kanR* | This study |
| SKY423 | *Mat a his3Δ1 leu2Δ0 ura3Δ0 met15∆0 RFC3-GFP::HisMX bar1::natR NUP170-3FLAG::kanR* | This study |
| SKY468 | *Mat a his3Δ1 leu2Δ0 ura3Δ0 met15∆0 CTF18-GFP::HisMX bar1::natR NUP60-3FLAG::kanR* | This study |
| SKY470 | *Mat a his3Δ1 leu2Δ0 ura3Δ0 met15∆0 CTF18-GFP::HisMX bar1::natR NUP192-3FLAG::kanR* | This study |
| SKY472 | *Mat a his3Δ1 leu2Δ0 ura3Δ0 met15∆0 CTF18-GFP::HisMX bar1::natR MLP1-3FLAG::kanR* | This study |
| SKY474 | *Mat a his3Δ1 leu2Δ0 ura3Δ0 met15∆0 CTF18-GFP::HisMX bar1::natR NUP2-3FLAG::kanR* | This study |
| SKY475 | *Mat a his3Δ1 leu2Δ0 ura3Δ0 met15∆0 CTF18-GFP::HisMX bar1::natR MLP2-3FLAG::kanR* | This study |
| SKY476 | *Mat a his3Δ1 leu2Δ0 ura3Δ0 met15∆0 CTF18-GFP::HisMX bar1::natR NUP84-3FLAG::kanR* | This study |
| SKY477 | *Mat a his3Δ1 leu2Δ0 ura3Δ0 met15∆0 CTF18-GFP::HisMX bar1::natR NUP120-3FLAG::kanR* | This study |
| SKY478 | *Mat a his3Δ1 leu2Δ0 ura3Δ0 met15∆0 CTF18-GFP::HisMX bar1::natR NUP133-3FLAG::kanR* | This study |
| SKY489 | *Mat a his3Δ1 leu2Δ0 ura3Δ0 met15∆0 CTF18-GFP::HisMX bar1::natR NUP1-6HA::kanR* | This study |
| SKY491 | *Mat a his3Δ1 leu2Δ0 ura3Δ0 met15∆0 CTF18-GFP::HisMX bar1::natR NUP53-3FLAG::kanR* | This study |
| SKY492 | *Mat a his3Δ1 leu2Δ0 ura3Δ0 met15∆0 CTF18-GFP::HisMX bar1::natR NUP59-3FLAG::kanR* | This study |
| SKY273 | *Mat a his3Δ1 leu2Δ0 ura3Δ0 met15∆0 NUP170-GFP1-10::URA3 NUP188-mCherry::hph RFC3-GFP11::natR* | This study |
| SKY281 | *Mat a his3Δ1 leu2Δ0 ura3Δ0 met15∆0 NUP170-GFP1-10::URA3 NUP188-mCherry::hph RFC2-GFP11::natR* | This study |
| SKY283 | *Mat a his3Δ1 leu2Δ0 ura3Δ0 met15∆0 NUP170-GFP1-10::URA3 NUP188-mCherry::hph CTF8-GFP11::natR* | This study |
| SKY310 | *Mat a his3Δ1 leu2Δ0 ura3Δ0 met15∆0 NUP170-GFP1-10::URA3 NUP188-mCherry::hph CTF18-GFP11::natR* | This study |
| SKY312 | *Mat a his3Δ1 leu2Δ0 ura3Δ0 met15∆0 NUP170-GFP1-10::URA3 NUP188-mCherry::hph RFC4-GFP11::natR* | This study |
| SKY314 | *Mat a his3Δ1 leu2Δ0 ura3Δ0 met15∆0 NUP170-GFP1-10::URA3 NUP188-mCherry::hph RFC5-GFP11::natR* | This study |
| SKY316 | *Mat a his3Δ1 leu2Δ0 ura3Δ0 met15∆0 NUP170-GFP1-10::URA3 NUP188-mCherry::hph DCC1-GFP11::natR* | This study |
| SKY346 | *Mat a his3Δ1 leu2Δ0 ura3Δ0 met15∆0 NUP49-GFP1-10::URA3 NUP188-mCherry::hph RFC3-GFP11::natR* | This study |
| SKY139 | *Matα his3Δ1 leu2Δ0 lys2Δ0 ura3Δ0 nup170∆::kanR* | (Giaever et al., 2002) |
| SKY530 | *Mat a his3Δ1 leu2Δ0 ura3Δ0 met15∆0 CTF18-GFP::HisMX bar1::natR elg1∆::zeoR* | This study |
| SKY531 | *Mat a his3Δ1 leu2Δ0 ura3Δ0 met15∆0 CTF18-GFP::HisMX nup170::kanR bar1::natR elg1∆::zeoR* | This study |
| SKY533 | *Mat α his3Δ1 leu2Δ0 lys2Δ0 ura3Δ0 ctf18∆::kanR* | (Giaever et al., 2002) |
| SKY534 | *Mat α his3Δ1 leu2Δ0 lys2Δ0 ura3Δ0 ctf8∆::kanR* | (Giaever et al., 2002) |
| SKY535 | *Mat α his3Δ1 leu2Δ0 lys2Δ0 ura3Δ0 dcc1∆::kanR* | (Giaever et al., 2002) |
| SKY484 | *Matα his3Δ1 leu2Δ0 lys2Δ0 ura3Δ0 nup170Δ::kanR elg1Δ::natR* | This study |
