## Supplementary material for "The nucleoporin Nup170 mediates subtelomeric gene silencing through the Ctf18-RFC complex and PCNA": Table S3

**RT-qPCR primers:**

| *COS4*fw | CCGTCATACCACTATGCTCTATTC |
| --- | --- |
| *COS4*rv | GCGTCAGTGTGCGATTCTAT |
| *COS10*fw | GCGTCGCCTAGATCATTAAAC |
| *COS10*fw | TGGTAACGGCTCATAGTTACA |
| *GIT1*fw | AGAGGTGGTATCCTGGTTATG |
| *GIT1*rv | CCTCCAGATCGCCTCTAAAT |
| *TOG1*fw | GTGAGCTCTGGCAATGTTAAT |
| *TOG1*rv | GATGCGTGTTAGGATGAAAGAA |
| *RFC1*fw | GCAGAGCTGTCCTATACGATTTA |
| *RFC1*rv | GGGTTTCCACATCCAACAAAG |
