## Supplemental figures for "The nucleoporin Nup170 mediates subtelomeric gene silencing through the Ctf18-RFC complex and PCNA"

**Figure S1 (related to Figure 1)**

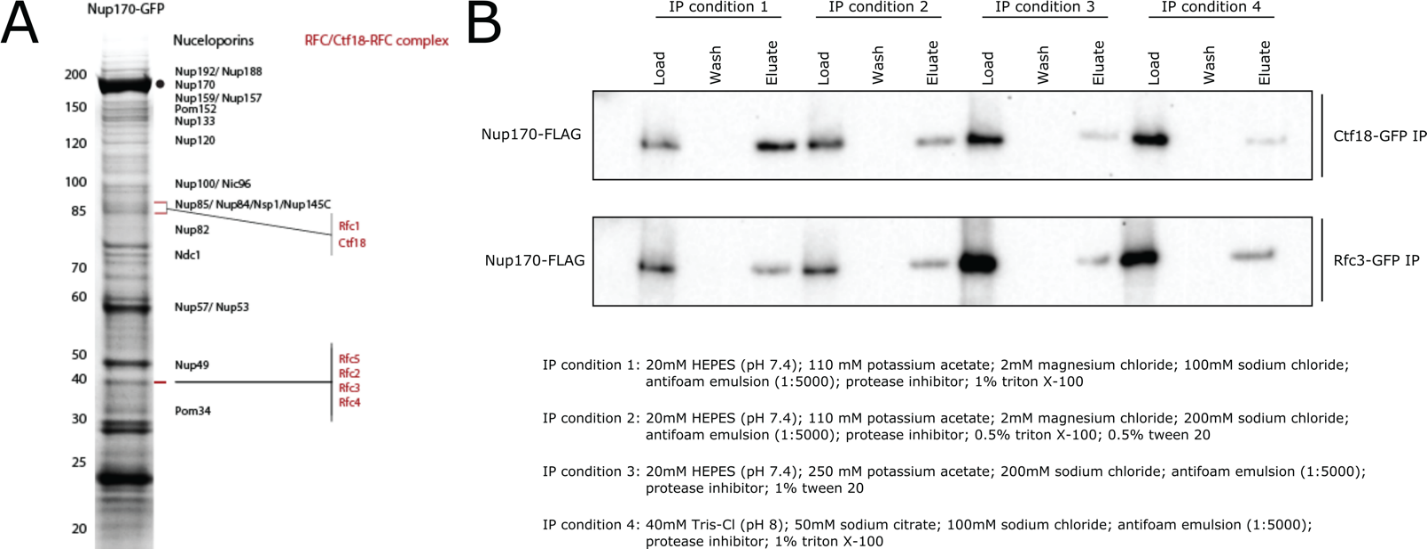

**Fig. S1. (**A) Nup170-GFP was affinity purified using a GFP nanobody and copurifying proteins were resolved by SDS-PAGE and visualized by staining with Coomassie blue. The gel bands were cut, processed by in-gel digestion, and analyzed by mass spectrometry. Proteins identified by mass spectrometry are listed in Table S1. (B) Ctf18-GFP and Rfc3-GFP fusion proteins were affinity purified from the cell lysate containing Nup170-3FLAG under various extraction conditions. Eluates were analyzed by immunoblotting using anti-FLAG antibody to detect co-immunoprecipitating Nup170.

**Figure S2 (related to Figure 4)**

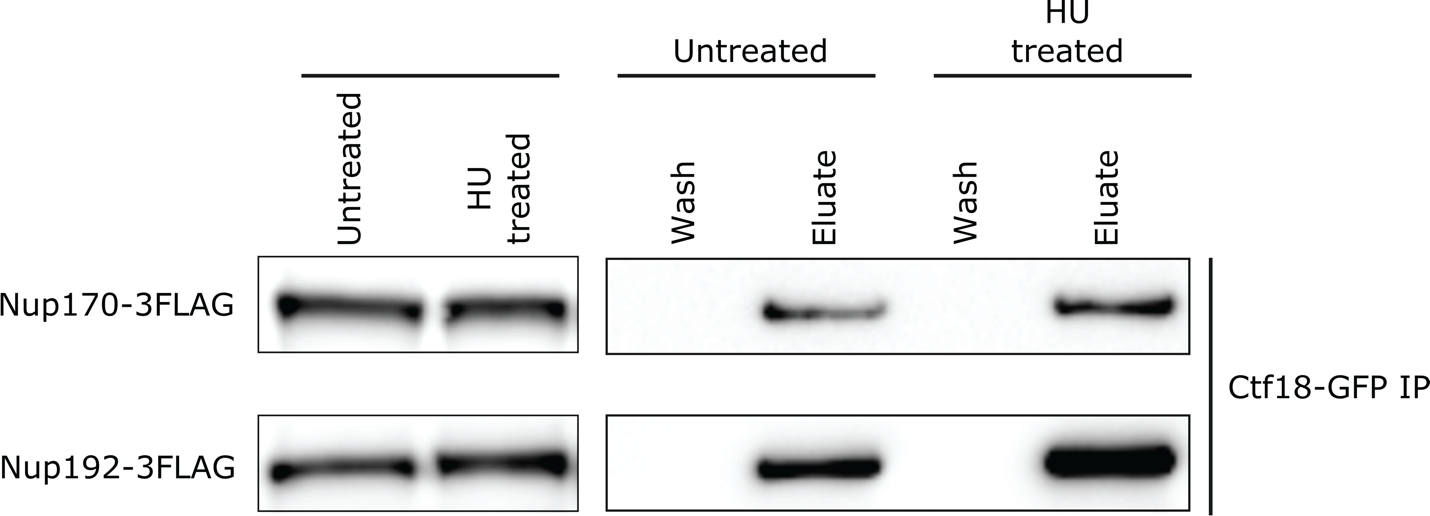

**Fig. S2**. **The NPC and Ctf18 binding increases after HU treatment**. Asynchronously growing cultures were treated with 200 mM HU for 100 min. The Ctf18-GFP fusion protein was affinity purified from cell lysates containing 3FLAG-tagged Nup170 or Nup192. Eluates were analyzed by immunoblotting using anti-FLAG antibodies to detect indicated nucleoporins.

**Figure S3 (related to Figure 5)**

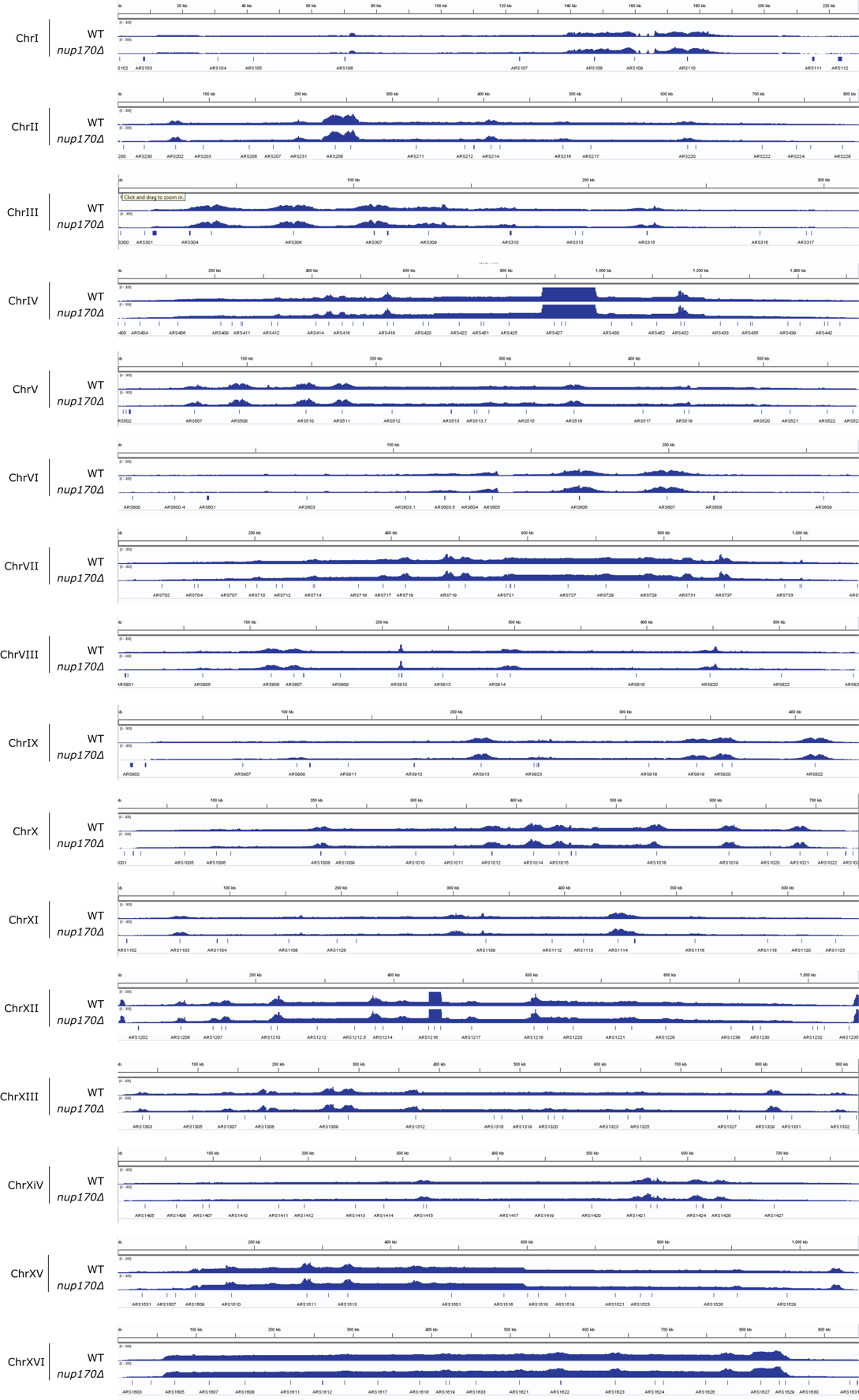

**Fig. S3.** **PCNA binding to all chromosomes**. Cells were synchronized in G1 and released into HU-containing YPD medium to arrest them in S phase, and ChIP was performed using an anti-PCNA antibody. Normalized PCNA binding profiles along chromosomes obtained by ChIP-seq analysis visualized with the Integrative Genomics Viewer.

**Figure S4 (related to Figure 6)**

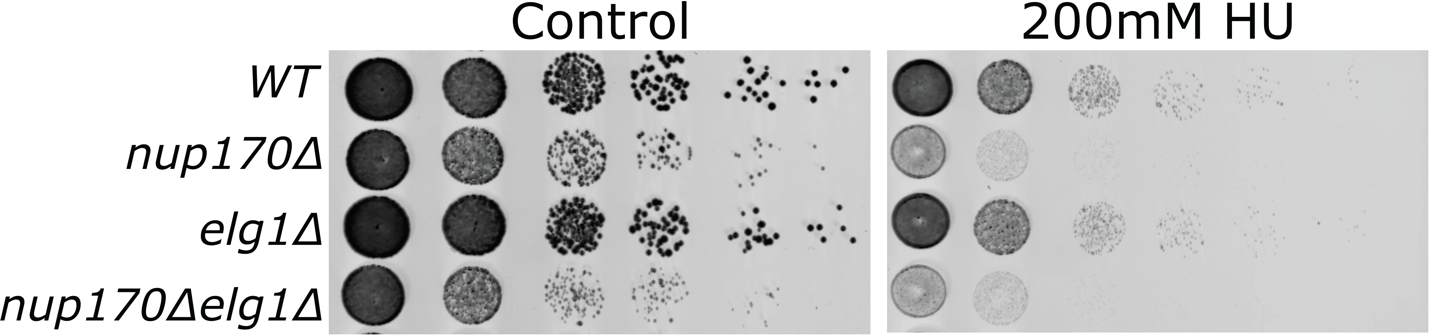

**Fig S4.** Log-phase cultures of the indicated strains were equalized in cellular density, serially diluted 10-fold, and spotted onto plates containing yeast extract, peptone, and dextrose (YPD) with or without 100 mM Hydroxyurea. Plates were scanned after 2 days at 30 °C.
